# Immunization of cows with HIV envelope trimers generates broadly neutralizing antibodies to the V2-apex from the ultralong CDRH3 repertoire

**DOI:** 10.1101/2024.02.13.580058

**Authors:** Pilar X. Altman, Gabriel Ozorowski, Robyn L. Stanfield, Jeremy Haakenson, Michael Appel, Mara Parren, Wen-Hsin Lee, Huldah Sang, Jordan Woehl, Karen Saye-Francisco, Collin Joyce, Ge Song, Katelyn Porter, Elise Landais, Raiees Andrabi, Ian A. Wilson, Andrew B. Ward, Waithaka Mwangi, Vaughn V. Smider, Dennis R. Burton, Devin Sok

## Abstract

The generation of broadly neutralizing antibodies (bnAbs) to specific HIV epitopes of the HIV Envelope (Env) is one of the cornerstones of HIV vaccine research. The current animal models we use have been unable to reliable produce a broadly neutralizing antibody response, with the exception of cows. Cows have rapidly and reliably produced a CD4 binding site response by homologous prime and boosting with a native-like Env trimer. In small animal models other engineered immunogens previously have been able to focus antibody responses to the bnAb V2-apex region of Env. Here, we immunized two groups of cows (n=4) with two regiments of V2-apex focusing immunogens to investigate whether antibody responses could be directed to the V2-apex on Env. Group 1 were immunized with chimpanzee simian immunodeficiency virus (SIV)-Env trimer that shares its V2-apex with HIV, followed by immunization with C108, a V2-apex focusing immunogen, and finally boosted with a cross-clade native-like trimer cocktail. Group 2 were immunized with HIV C108 Env trimer followed by the same HIV trimer cocktail as Group 1. Longitudinal serum analysis showed that one cow in each group developed serum neutralizing antibody responses to the V2-apex. Eight and 11 bnAbs were isolated from Group 1 and Group 2 cows respectively. The best bnAbs had both medium breadth and potency. Potent and broad responses developed later than previous CD4bs cow bnAbs and required several different immunogens. All isolated bnAbs were derived from the ultralong CDRH3 repertoire. The finding that cow antibodies can target multiple broadly neutralizing epitopes on the HIV surface reveals important insight into the generation of immunogens and testing in the cow animal model. The exclusive isolation of ultralong CDRH3 bnAbs, despite only comprising a small percent of the cow repertoire, suggests these antibodies outcompete the long and short CDRH3 antibodies during the bnAb response.

**Author Summary:** The elicitation of epitope-specific broadly neutralizing antibodies is highly desirable for an HIV vaccine as bnAbs can prevent HIV infection in robust animal challenge models and humans, but to date, cows are the only model shown to reliably produce HIV bnAb responses on Envelope (Env) immunization. These responses involve Abs with ultralong CDRH3s and are all directed to a single site, the CD4 binding site. To determine whether this is a unique phenomenon or whether cow antibodies can target further bnAb sites on Env, we employed an immunization protocol that generated cow bnAbs to a second site, the V2-apex. We conclude that ultralong CDRH3s are well adapted to penetrate the glycan shield of HIV Env and recognize conserved regions and may constitute protein units, either in the context of antibodies or in other engineered proteins, that could be deployed as anti-HIV reagents.

## Introduction

Broadly neutralizing antibodies (bnAbs) neutralize diverse HIV isolates by recognizing relatively conserved epitopes on the HIV Env trimer, and the elicitation of such antibodies by vaccination is widely considered a key component of an efficacious vaccine. Cohort studies have proven that humans infected with HIV are capable of developing bnAbs, although the antibodies typically have unusual features such as high levels of somatic hypermutation and longer-than-average loops in the complementarity determining region of the heavy chains (CDRH3)[1]. Animals immunized with recombinant Env have been shown to have immune responses overwhelmingly to non-neutralizing epitopes[2-6]. Models have been put forth to explain these results, including affinity disparity model and cell number disparity models[5]. The affinity disparity model suggests that B cells targeting non-neutralizing epitopes have higher affinities for antigen and out-compete neutralizing responses that have comparatively lower affinities. The cell number disparity model proposes that the frequencies of naïve B cells with specificities to neutralizing epitopes are rare relative to naive B cells targeting non-neutralizing epitopes and are therefore also outcompeted in germinal centers. Both models focus on the idea that an immunodominant response to certain epitopes overwhelms the development and affinity maturation of more immunoquiescent responses such as those to bnAb epitopes. To overcome this problem, immunogens will need to properly prime and expand naïve B cell responses to neutralizing epitopes taking into consideration precursor frequencies and affinities.

The V2-apex is a promising epitope on HIV Env for the development of a vaccine designed to elicit bnAbs. Appropriate immunogens can be designed to select for rare B cells with B cell receptors (BCRs) that are capable of penetrating through the glycan shield at the trimer apex including two prominent glycans at position N156 and N160[7, 8]. The bnAbs that target this epitope region typically have very long CDRH3 loops, extending up to 37 amino acids, with tyrosine sulfate post-translational modifications and negatively charged amino acids that enable or enhance binding to the positively charged C-strand of the V2 loop on the Env trimer[9-12]. Such antibodies are rare in the naïve human antibody repertoire where CDRH3 of > 24 amino acids (AAs) and > 28 AAs have frequencies of 3.5% and 0.43%, respectively[13].These long CDRH3 antibodies are similarly rare in preclinical animal models such as rabbits and nonhuman primates (NHPs) and their elicitation presents a major challenge for immunogen design strategies[14-17]. Given their capacity to produce exceptionally long CDRH3s, cows are an interesting model system for evaluating immunogens designed to elicit V2-apex bnAbs. The cow antibody repertoire contains a unique subset (∼10%) of antibodies possessing ultralong CDRH3s, which can reach lengths between 50 to 70 amino acids[18-22]. In addition, the cow antibody repertoire is skewed toward longer CDRH3s with an average length of 26 amino acids as compared to 15 in the human repertoire[14, 18, 22-30]. We define here “ultralong” CDRH3 cow antibodies as those having CDRH3 lengths of ≥50 amino acids and “long” CDRH3 cow antibodies as those having lengths of 25-49 amino acids. Cow antibodies having CDRH3 lengths <25 amino acids are defined as “short”. The ultralong cow CDRH3s are all encoded by the same V and D gene segments, IGHV1-7 and IGHD8-2 respectively[18, 19, 31, 32]. The CDRH3 length is primarily encoded by the IGHD8-2 germline gene segment which encodes 48-50 amino acid residues depending on the polymorphic variant[31-33]. The unique CDRH3 structure can be separated into two microdomains where the antigen-binding disulfide-bound “knob” sits on top of a long b-ribbon “stalk” and can bind to both surface and receded epitopes[22, 34, 35]. As cows have lower VDJ combinatorial diversity potential compared to humans, they must utilize different strategies to create diversity and length in their repertoires[22, 36]. Some of their diversity is due to V-D and D-J junctions which can alter the amino acid length and therefore the stalk of the ultralong CDRH3[33, 34, 36-38]. The knob, which appears to be the most important region for antigen interaction, requires somatic hypermutation for diversity. The somatic hypermutation occurs both before and after antigen exposure and introduces diversity into the knob using amino acid point mutations, nucleotide deletions, and mutations that substitute to and from cysteine[18, 22, 33, 38-40]. The knob residues encoded by IGHD8-2 include four conserved cysteines as well as repeated glycine, tyrosine, and a few serine amino acid residues. These residues have a severe codon bias towards mutation to cysteine, which can introduce additional diversity in structure in the form of disulfide bond formation[22, 33, 35, 41].

Previous studies on immunizations of cows with non-well-ordered HIV Env trimers resulted in some neutralization breadth and potency against cross-clade HIV pseudoviruses in colostrum[42]. The antibody activity from these cows was CD4bs specific[42-45]. An immunization study of four cows with a recombinant, well-ordered trimer, BG505 SOSIP.664, elicited broad and potent serum antibody responses in as little as 42 days[46, 47]. Strikingly, the responses were also CD4bs specific with no evidence of responses targeting any other bnAb epitopes, including the V2-apex epitope. The monoclonal antibodies isolated were determined to be broadly neutralizing, had ultralong CDRH3 regions, and are some of the most potent bnAbs to the CD4bs isolated to date[47]. A more recent study even employed the use of cross-clade trimers and heterologous baits for sorting. While isolated antibodies from this study were extremely potent and broad, they also targeted the CD4bs[48].

To determine whether cow antibodies to another bnAb epitope could be elicited given the ultralong CDRH3s, we used a V2-apex focusing immunogen strategy. In the study, two groups of two cows were immunized over one year with different immunization regimens; the immunogens used were selected based on their sensitivity to V1V2 bnAb inferred precursors[2, 7, 8]. Cows in Group 1 were primed and boosted twice with a stabilized native-like chimpanzee SIV trimer MT145K SOSIP followed by two boosts of a stabilized native-like HIV trimer, C108 SOSIP, and finally boosted with a cocktail of stabilized near-native HIV SOSIP trimers. The rationale for the SIV prime/HIV Env boost was that the V2-apex bnAb site was the only epitope largely conserved between SIV and HIV for the chosen isolates, so we aimed to focus responses to this site using a prime-boost strategy. The cocktail immunization was added as a final step to increase the breadth of nAb responses. Cows in Group 2 were primed and boosted four times with C108 SOSIP followed by one boost with the SOSIP cocktail used in Group 1. In both groups, the recombinant SOSIP cocktail consisted of trimers derived from isolates CRF250 (clade AE), WITO (clade B), ZMZM (ZM197 backbone with ZM233’s V1V2 region, clade C), and BG505 (clade A). Sera were collected longitudinally from these animals to monitor the development of bnAb responses, and IgG^+^ B cells were subsequently sorted and sequenced from peripheral blood mononuclear cells (PBMC) or splenocytes of cows that showed broadly neutralizing activity. Monoclonal antibodies were recombinantly produced from memory B cells and further characterized.

## Results

### Cows immunized with V2-apex focusing immunogens elicit a delayed cross-clade neutralizing response compared to BG505 SOSIP

Group 1, comprising cow-485 and cow-16157, were primed with SIV MT145K SOSIP on days 37 and 79, boosted with HIV C108 SOSIP on days 142 and 226, and finally boosted on day 336 with a cocktail of recombinant HIV SOSIP trimers. Group 2, comprising cow-488 and cow-491, were primed and boosted with HIV C108 SOSIP and finally boosted with the same cocktail of recombinant HIV SOSIP trimers[49, 50]. The immunization protocols are detailed in Fig 1A. Following completion of the immunization, IgGs from terminal bleed sera (day 361 post prime) were purified and tested for their ability to neutralize autologous viruses (Fig 1B). Cows from both groups developed autologous neutralizing responses to some but not all of the immunogens. Cow-485 developed neutralization activity against all autologous viruses except WITO and had the second-highest autologous titers overall, while cow-16157 had moderate breadth and potency but was eliminated from further evaluation due to nonspecific neutralizing activity to murine leukemia virus (MLV) for some samples (data not shown). Cow-488 developed neutralization to all autologous viruses tested, but cow-491 from the same group had the least broad and potent autologous neutralizing response.

**Fig 1:**
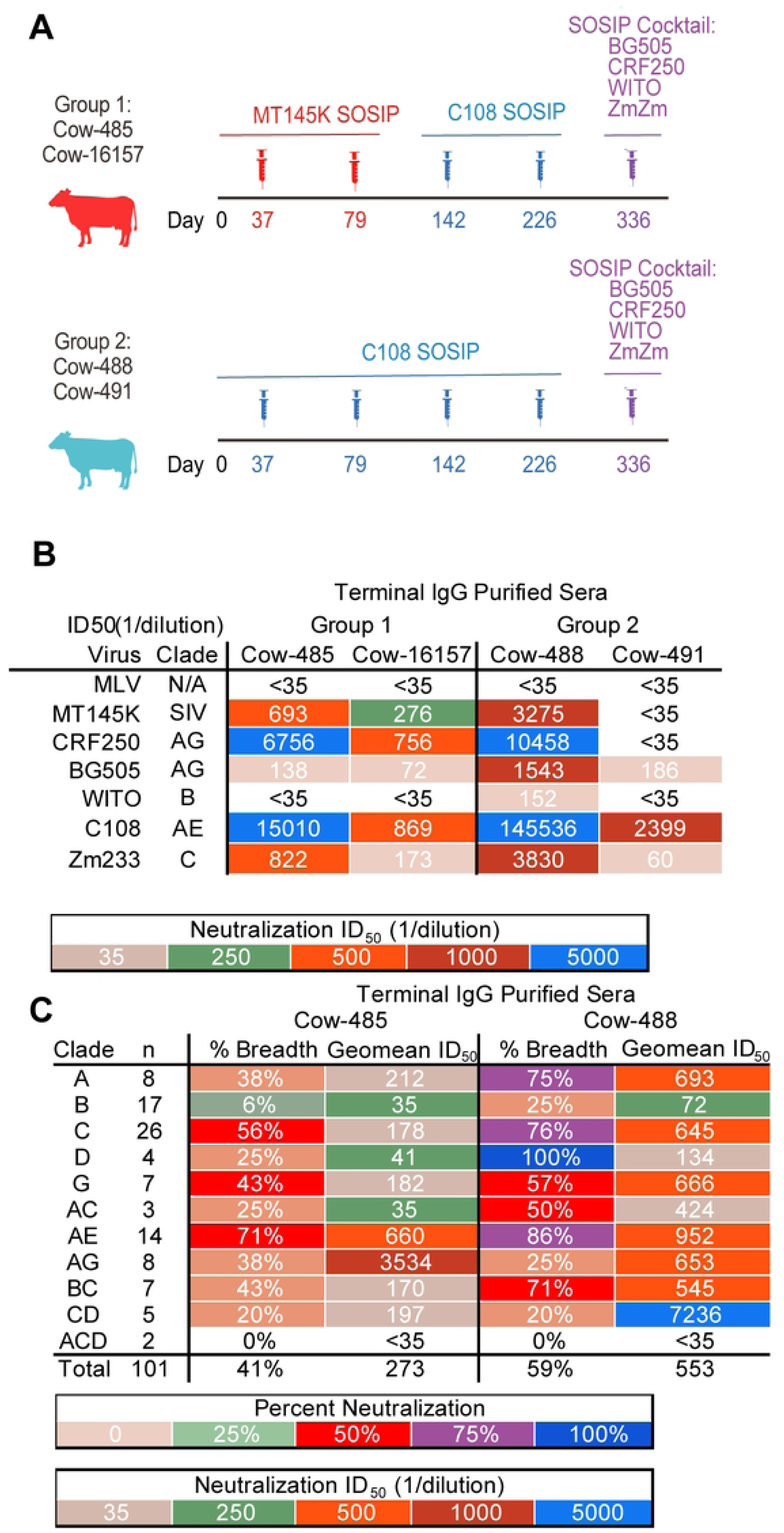
Cows were immunized with a series of V2-apex focusing immunogens. **(A)** Schematic of SOSIP immunization experiments in two groups of cows. Group 1 is shown in red, Group 2 is shown in light blue. Syringes indicate the timing of boosts. Immunizations with MT145K SOSIP are shown in red, C108 SOSIP in blue, and the SOSIP cocktail in purple. **(B)** Terminal serum nAbs ID_50_ titers were tested for neutralization for all four cows against the autologous viruses. MLV is a negative control. **(C)** Terminal IgG-purified sera were tested for neutralization by sera from cow-488 and cow-485 on a large 101 virus cross-clade panel. Neutralization breadth and geomean ID_50_ titers are grouped by virus clade.

Day 359 was evaluated for neutralization on a 12-virus global panel for heterologous neutralization (S1 Table)[51]. All cows were able to neutralize heterologous viruses from the panel. Cow-488 from Group 2 was the most broad and potent followed by cow-485 from Group 1. Based on their cross-clade neutralization activities, cow-485 and cow-488 were selected for further evaluation. To assess the neutralization breadth and potency for cow-485 and cow-488, purified serum IgGs from the terminal time point were further evaluated for neutralization activity on a 101 cross-clade pseudovirus panel[52] (Fig 1C). Cow-485 and cow-488 neutralized 41% and 59% of viruses, respectively. Cow-485 poorly neutralized clade B and clade ACD with only 6 and 0% of the viruses neutralized respectively. While cow-488 had better neutralization breadth overall, the two clade ACD viruses were still not neutralized and clade B, AG, and CD were only neutralized 20-25% respectively.

We next evaluated the development of the cross-clade longitudinal neutralization and selected 13 time points to test neutralization on the 12-virus global panel (Fig 2). Cow-485’s cross-clade clade neutralization mostly developed after the addition of a SOSIP cocktail, although clade A virus CNE55 developed neutralization before the cocktail. Three of the four viruses that were not neutralized were clade B or clade BC. Cow-488 neutralized four of the 12 viruses before addition of the cocktail. Once the SOSIP cocktail was used for immunization, the response broadened across the panel. Overall, all the cows displayed some level of autologous neutralization and cow-485 and cow-488, in particular, displayed broad cross-clade neutralization following boosting with a cocktail of immunogens.

**Fig 2:**
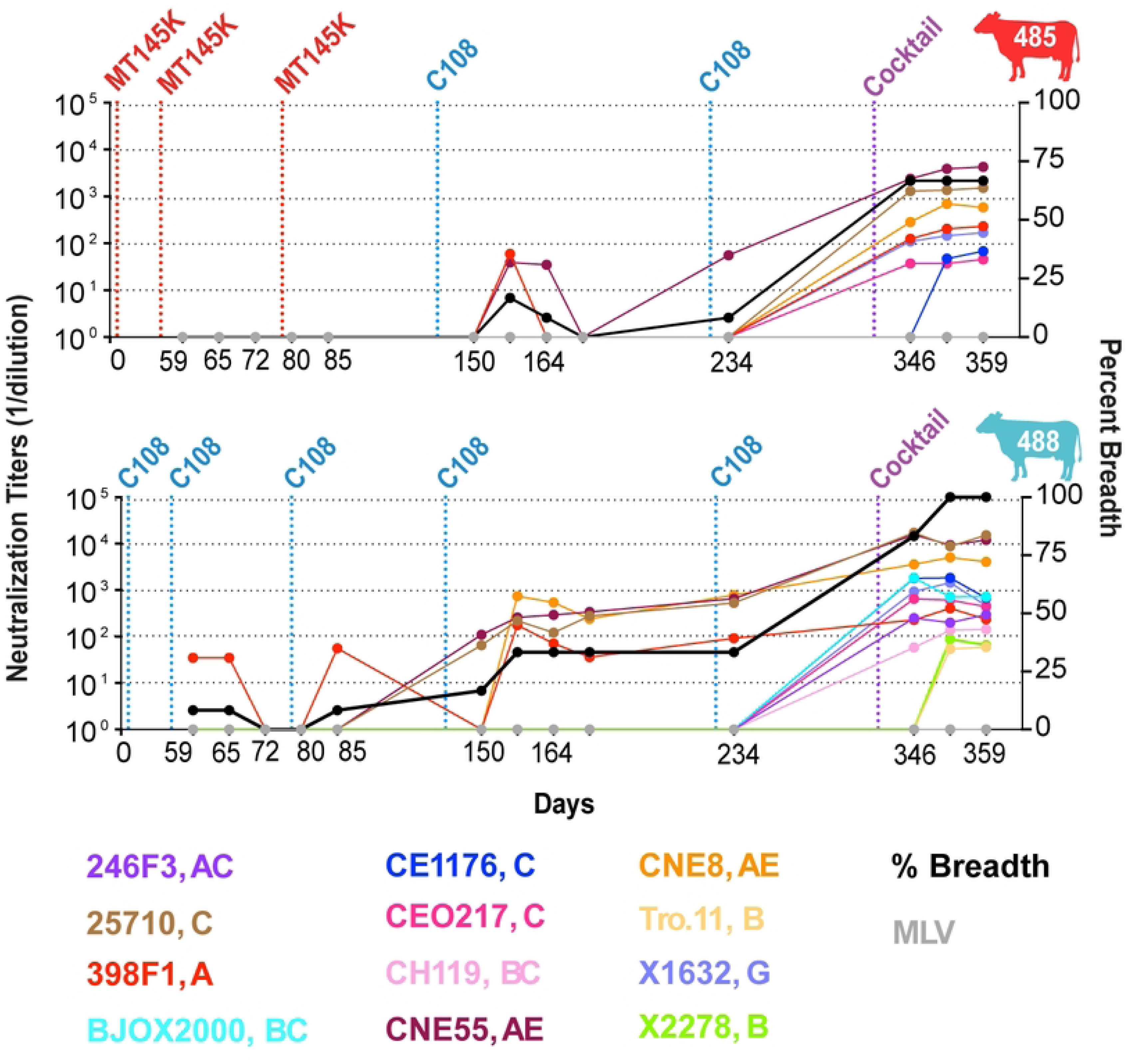
Cows develop serum broadly neutralizing antibody responses following immunization with recombinant trimers. Longitudinal samples were collected and tested for cow-485 and cow-488 on the 12-virus global panel. Neutralization ID_50_ titers are presented for each virus by color for each time point. Vertical dotted lines represent prime and boost dates, immunogen colors are described in Fig 1A. Neutralization percent breadth is represented by a black line whose y-axis is shown on the right. ID_50_ values are shown as 1/dilution on the y-axis.

### Epitope mapping of sera from cow-485 and cow-488

After confirming the development of broad and potent neutralizing serum responses in two cows, we next attempted to map the epitope specificities in sera. We first used polyclonal electron microscopy epitope mapping (EMPEM) of Fabs digested from IgGs purified from sera at the terminal time point and binding to BG505 SOSIP. EMPEM revealed Fab densities for both cows at a neoepitope at the base of the trimers, consistent with responses for other animals (data not shown). Cow-485 showed some V2-apex response in a single 2D class but it could not be reconstructed (data not shown). We next performed competition ELISAs using cow sera from multiple time points and biotinylated HIV bnAbs to assess polyclonal epitope specificities on BG505 SOSIP (Fig 3A). IgGs purified from cow-485 and cow-488 sera from day 85, day 157, day 234, and day 352 were tested for competition against mAbs targeting the V2-apex including CAP256-VRC26.9 and PGDM1400, CD4bs targeting NC-Cow1, gp120/gp140 interface targeting 35O22, and V3 glycan directed PGT121. Cow-488 had more than 50% competition with apex bnAbs CAP256-VRC26.9 and PGDM1400 on day 346 and day 352, in both cases after the SOSIP cocktail immunization. Cow-485 sera did not show more than 50% competition with the bnAbs tested.

**Fig 3:**
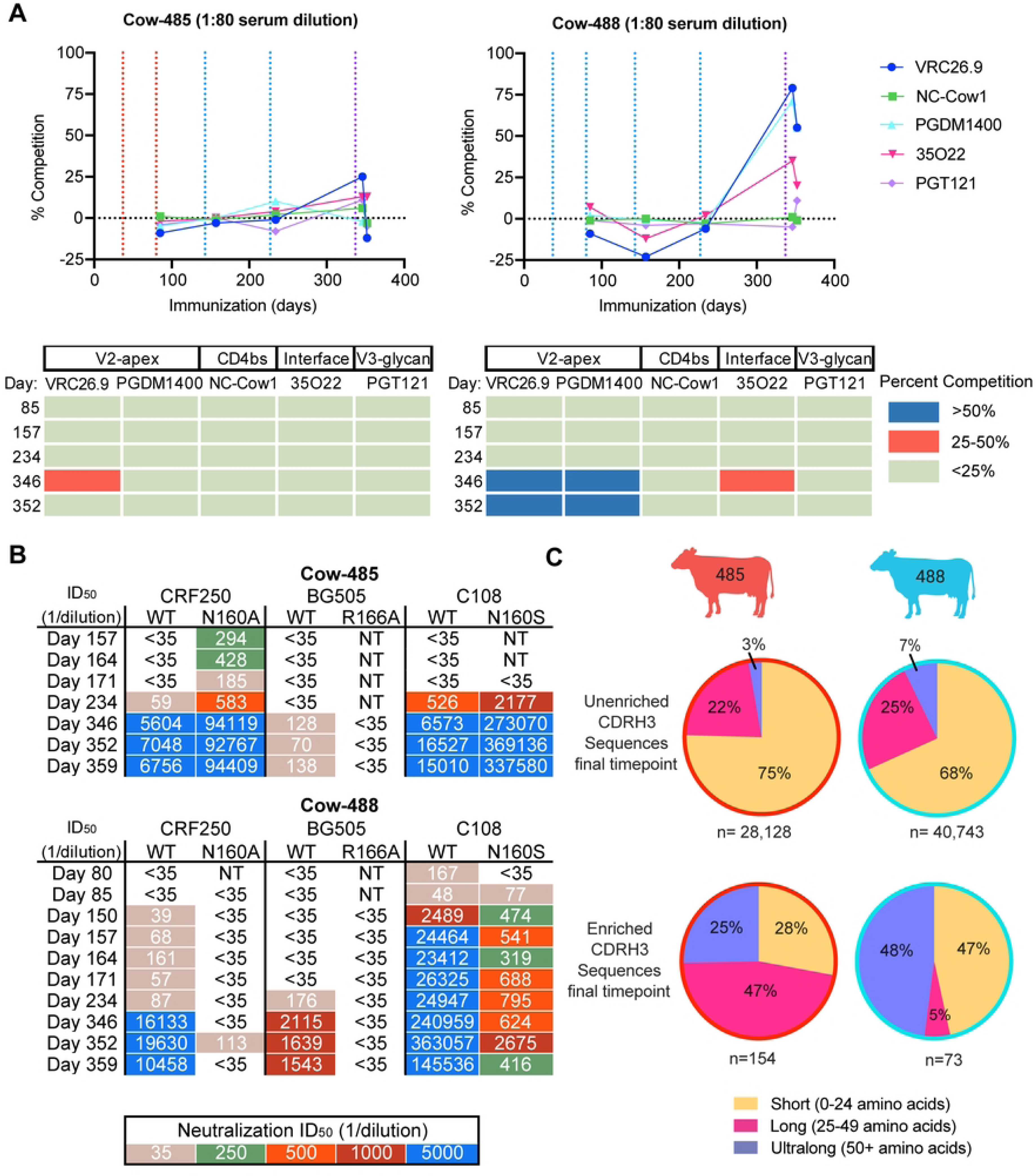
Broadly neutralizing responses in immunized cows are directed to the V2-apex epitope on HIV Env. **(A)** Sera from cow-485 and cow-488 over five time points were tested for competition against five well characterized antibodies spanning four HIV bnAb epitopes. Vertical dotted lines represent prime and boost dates. Immunizations of MT145K SOSIP are shown in red, C108 SOSIP in blue, and the SOSIP cocktail in purple. The results are displayed both graphically and tabulated percent neutralization. CAP256-VRC26.9 is referred to as VRC26.9. **(B)** Neutralization ID_50_ titers are shown for IgG-purified sera tested for neutralization of wild-type and V2-apex epitope mutant viruses spanning multiple time points. NT= not tested. **(C)** Comparisons of the amino acid CDRH3 lengths for unenriched and sort-enriched (see text) terminal samples from cow-485 and cow-488 are shown in a pie chart. Percentage of whole are indicated in each section of the pie. Short CDRH3 length antibodies are shown in yellow, long in pink, and ultralong in purple.

Importantly, the EMPEM and competition ELISAs provide an indicator of immunodominant binding responses. We therefore next determined if we could measure neutralizing activity directed to the V2-apex. First, we evaluated neutralizing activity against HIV pseudoviruses CRF250, BG505, and C108 wild-type (WT) viruses as well as the same viruses with variants containing substitutions at amino-acid residues R166 and N160, which are all localized at the trimer apex epitope (Fig 3B). Serum neutralizing activity was abrogated for BG505 R166A but not CRF250 N160A or C108 N160S mutants in cow-485. Neutralization titers increased for the N160 mutants and there was neutralization of CRF250 N160A at time points for which the WT virus was not neutralized, suggesting that the glycan at position N160 may be inhibiting the polyclonal neutralizing response for cow-485 from surrounding regions. For cow-488, the neutralizing serum response was similarly abrogated in BG505 R166A and CRF250 N160A variants. Neutralizing titers were reduced for C108 N160S variant virus in cow-488 as early as day 80, suggesting important residues in the V2-apex were targeted by neutralizing antibodies in the sera early in the immunization series. Both cows appear to have some level of V2-apex targeting neutralizing antibody responses.

### Broadly neutralizing antibodies isolated from cow-485 and cow-488 have ultralong CDRH3s and medium neutralization breadth and potency

After confirming V2-apex-directed nAb responses, we next attempted to isolate monoclonal antibodies from cow-485 and cow-488 using four rounds of single IgG^+^ B cell sorting. We used PBMCs and splenocytes from several late time points from both cows. Sorts 1 and 2 used the terminal time point for both cow-485 and cow-488. Sorts 3 and 4 used PBMC/splenocyte samples from day 352 and day 361 for both cow-485 and cow-48. All sorts were stained with goat anti-cow IgG conjugated with FITC and biotinylated antigens conjugated to streptavidin fluorophores to isolate the B-cell population of interest. All SOSIP baits were biotinylated and stained with streptavidin conjugated to fluorophores. Sort 1 and Sort 2 were done the same with only higher affinity cell sorted in Sort 2[53] (S1 Fig.). The third sort selected for cells that demonstrated double positive binding to the same SOSIP (MT145K or 25710) coupled individually to PE or AF-647 to isolate high affinity cells and eliminate those with non-specific binding. Sort 4 isolated cells positive to binding only MT145K SOSIP (PE) and did not have affinity for MT145K-PGT145 Fab Complex (AF-647) to isolate cells targeting the V2-apex. Variable regions of isolated B cells were recovered using single-cell PCR amplification and subsequently cloned into human antibody expression vectors as previously described[47]. Cow-485 had a much higher heavy chain recovery in the first three sorts and therefore had more antibodies tested overall (S2 Table). The unenriched B cell repertoires were sequenced for cow-485 and cow-488 at the terminal time point and compared to the sequences recovered from the repertoires isolated during the four sorts (Fig 3C). Cow-485’s enriched response was heavily weighted toward long CDRH3 antibodies, while cow-488 had a small percentage of long CDRH3 antibodies and a large percent of ultralong and short CDRH3 antibodies in the enriched pool. As a whole, the antibodies isolated during the sorts revealed an enrichment of long and ultralong CDRH3 antibodies in trimer-immunogen specific B-cells compared to the total repertoire.

We then examined the CDRH3 sequences of isolated antibodies with different lengths to identify tyrosine sulfation motifs, which are typically important for neutralization by human V2-apex bnAbs. Although tyrosine sulfation is challenging to predict, we focused on the DY and YD motifs for this study. Ultralong antibodies had the highest number of DY/YD motifs in total and as a percentage of their overall population; 60% of the ultralong CDRH3 antibodies in cow-485 and 80% in cow-488 had the motif (Fig 4A). We recombinantly expressed all isolated antibodies with their native light chain and/or a universal cow light chain V30 (termed a ‘universal’ cow light chain), also denoted Vlx1, that is predominantly paired with ultralong CDRH3 heavy chains and subsequently evaluated them for neutralization against heterologous viruses[27] (Fig 4B, S2 Table). IGHV1-7 derived antibodies from the first three sorts were also tested with the universal cow light chain. We included the universal cow light chain to assure all heavy chains recovered were tested as thoroughly as possible. While we did test short and long IGHV1-7 derived antibodies with the universal cow light chain, they do not always pair with this light chain, but were tested and included in the results regardless. We screened antibodies using high-throughput expression of antibodies in Expi293 cells and tested the supernatants for expression and binding to BG505 SOSIP using ELISA. Antibodies were tested with their native light chain when both heavy chain and light chain were isolated for all four sorts. The supernatants of expressed antibodies were then screened for neutralization of C108, CRF250, and CNE55 viruses. For cow-485 a total of 134 heavy chains paired with native light chain and 80 heavy chains paired with universal light chain. Of those paired with the native light chains, 41 of the heavy chains had CDRH3s that were short, 63 were long, and 30 were ultralong. Of those paired with a universal light chain, none were short, 52 were long and 28 were ultralong. For cow-488 a total of 71 mAbs paired with their native light chain and 3 mAbs paired with universal light chains were tested. In the native light chain set, 33 of the heavy chain CDRH3s were short, 3 were long, and 35 were ultralong (S2 Table). Of those paired with a universal light chain, 1 of the heavy chain CDRH3s was short, 2 were long and none were ultralong. Ultimately, we isolated a total of eight and 11 cross-clade nAbs from cow-485 and cow-488, respectively, all of which had ultralong CDRH3s, suggesting these antibodies were responsible for cross-clade neutralization detected in sera.

**Fig 4:**
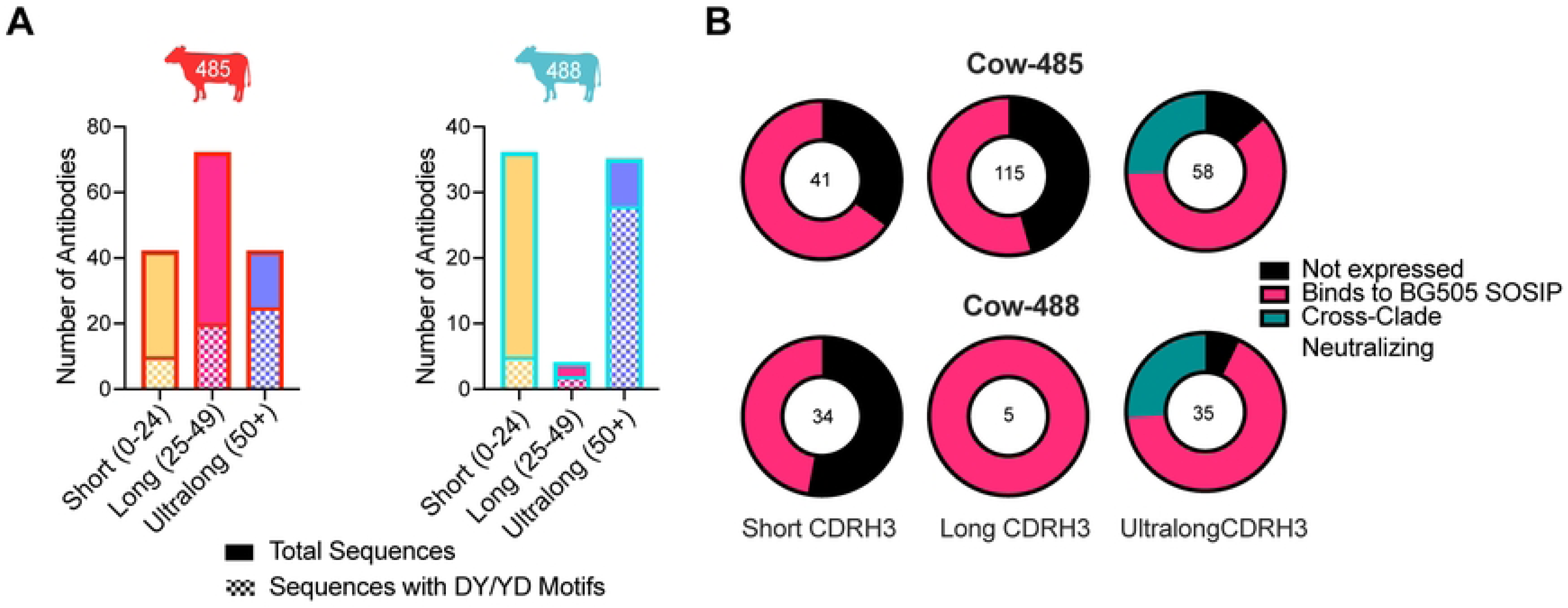
Broadly neutralizing antibodies are ultralong and have medium breadth and high potency. **(A)** Sort-enriched isolated CDRH3 lengths were grouped into bars to show the ratio of those with DY/YD motifs. Short CDRH3 length antibodies are shown in yellow, long in pink, and ultralong in purple. Bars are outlined in red for cow-485 and blue for cow-488. **(B)** Pie charts representing the total antibodies isolated and tested are shown and categorized by CDRH3 length. Colors indicate antibodies that did not express in black, those that bound to BG505 SOSIP using ELISA in pink, and those that were determined to have cross-clade neutralization in teal. The number in the middle of the graph represent the total isolates and tested antibodies. **(C)** CDRH3s are shown for Bess and ElsE antibodies. The sequences for the germline VH1–7, DH8-2, and JH10 regions are shown at the top. Number of cysteines (#Cys) and CDRH3 lengths (L) are shown to the left. Cysteines within the CDRH3s are highlighted in blue. Negatively charged D and E amino acids are shown in red. **(D)** Summary results of Bess and ElsE series mAb measured against a 12-virus global panel for neutralization. Neutralizing geomean IC_50_ and geomean IC_80_ titers are shown for viruses neutralized at IC_50_ <50 µg/ml. Percent breadth on the panel is shown for each as well as the geomean maximum percent neutralization (MNP) for viruses neutralized at IC_50_ <50 µg/ml. **(E)** MAbs Bess1, Bess2, Bess4, ElsE1, and ElsE2 were analyzed for neutralization breadth and potency on 101 viruses of a muti-clade panel. Neutralization breadth and geomean IC_50_ values are grouped by virus clade.

We next aligned the CDRH3 sequences of the isolated antibodies with the germline and bnAb NC-Cow1[22] (Fig 5A). We focused on cysteines responsible for the structure of ultralong CDRH3 antibodies and negatively charged amino acids in CDRH3. Antibodies from cow-485 were named Bess, while antibodies from cow-488 were named ElsE and assigned numbers based on their order of discovery. The Bess and ElsE antibodies, as for all ultralong cow antibodies, derive from the same germline VH, DH, and JH genes so there are expected similarities in overall sequence. When aligned as the full variable regions, the antibodies grouped together phylogenetically (S2 Fig.) and when aligned with only CDRH3s, the lineages did separate phylogenetically into different branches with some somatic variants as exceptions—Bess5 and Bess4 grouped phylogenetically with ElsEs mainly due to similarities in the region most likely to be the “stalk”. Both ElsE and Bess antibodies contained a significant number of negatively charged amino acids in the area likely to be the knob, based on another ultralong NC-Cow1 bnAb structure[35]. This feature is important for the ability of human bnAbs to bind to the positively charged lysine patch of the V2-apex. We tested all antibodies from the Bess and ElsE lineages for neutralization of viruses from the 12-virus global panel (Fig 5B, S3 Table). The broadest Bess antibodies were able to neutralize seven of the 12 viruses from the global panel, while the least broad and potent only neutralized three. The most broad and potent nAb of the ElsE lineage was able to neutralize eight, while the least potent neutralized five viruses. Antibodies from both lineages showed some incomplete neutralization (<95%).

**Fig 5:**
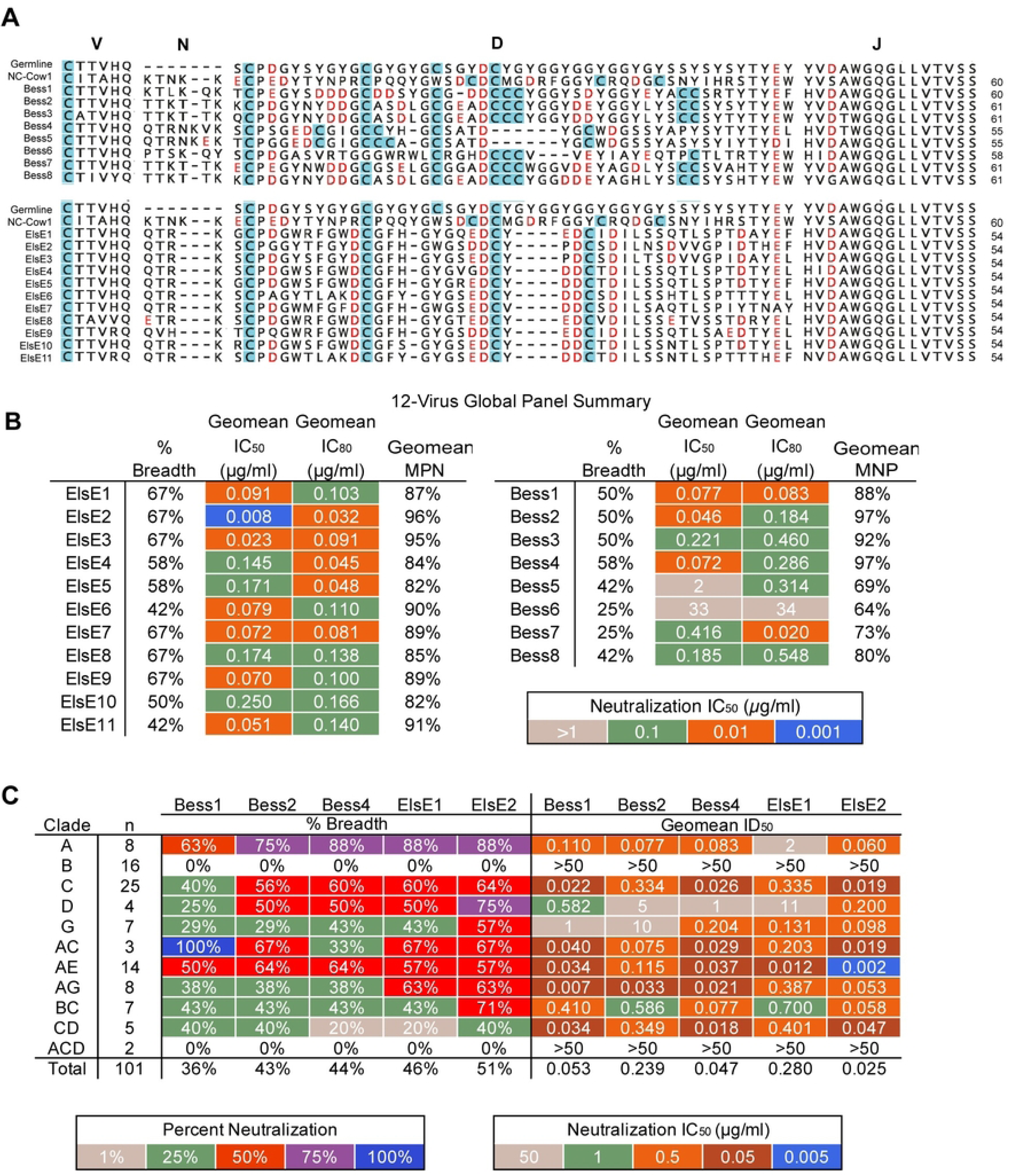
Bess and ElsE broadly neutralizing antibodies target the V2-apex trimer and not all require tyrosine sulfation for neutralization. **(A)** Bess4 and ElsE2 were mapped on isolate 25710 SOSIP using BLI epitope binning. Competing antibodies are categorized according to epitope. CAP256-VRC26.9 is referred to as VRC26.9. Percent competition is shown. **(B)** Bess and ElsE neutralizing IC_50_ titers were evaluated for neutralization of wild-type and V2-apex epitope-mutated residue viruses. **(C)** Bess 1 neutralization is reduced on elimination of a sulfation motif in CDRH3. The CDRH3 of Bess1_WT and the mutated Bess1_DF are shown together with comparative neutralization IC_50_ values in μg/ml on a bar chart.

We chose Bess1, Bess2, Bess4, ElsE1, and ElsE2 mAbs as representatives to be tested on 101 viruses from a large cross-clade panel[51, 52] (Fig 5C). The ElsE lineage was the most potent and broad overall. The neutralization patterns for both Bess and ElsE antibodies were similar to those of human V2-apex antibodies CAP256-VRC26, which were unable or poorly able to neutralize most Clade B viruses. However, ElsE and Bess antibodies were very broad and potent against Clade A viruses. Additionally, Bess and ElsE antibodies tested on the panel did show some incomplete neutralization with plateauing at <100% neutralization or non-sigmoidal curves. This behavior is not uncommon among trimer-specific antibodies[54-58]. Overall, the antibodies were moderately broad and potent, but less broad and less potent than the CD4bs-targeting cow bnAb NC-Cow1[47].

### Bess and ElsE broadly neutralizing antibodies target the V2-apex trimer and not all require tyrosine sulfation for neutralization

To assess epitope specificities of the recombinant mAbs, Bess4 and ElseE2 were selected as Bess and ElsE representatives for competition biolayer interferometry (BLI) against a panel of human antibodies that target known bnAb epitopes on the Env trimer of virus 25710 as well as NC-Cow1. The BLI experiments demonstrated competition of Bess and ElsE representatives with antibodies CAP256-VRC26.9 and PGT145 (Fig 6A). These results suggest Bess and ElsE are targeting the V2-apex. As V2-apex targeting antibodies are trimer specific or preferring, we used an ELISA binding assay to determine if Bess or ElsE antibodies could bind to monomeric gp120. Both lineages were unable to bind to the monomer (S3A Fig.). SPR data were generated to quantify binding affinity to BG505 trimers, which ranged in K_D_ from 858 to 17 nM. Not all of the antibodies bound to BG505, although the potent bnAbs did have highest affinity reaching a K_D_ of 17 nM (S4 Fig.).

**Fig 6:**
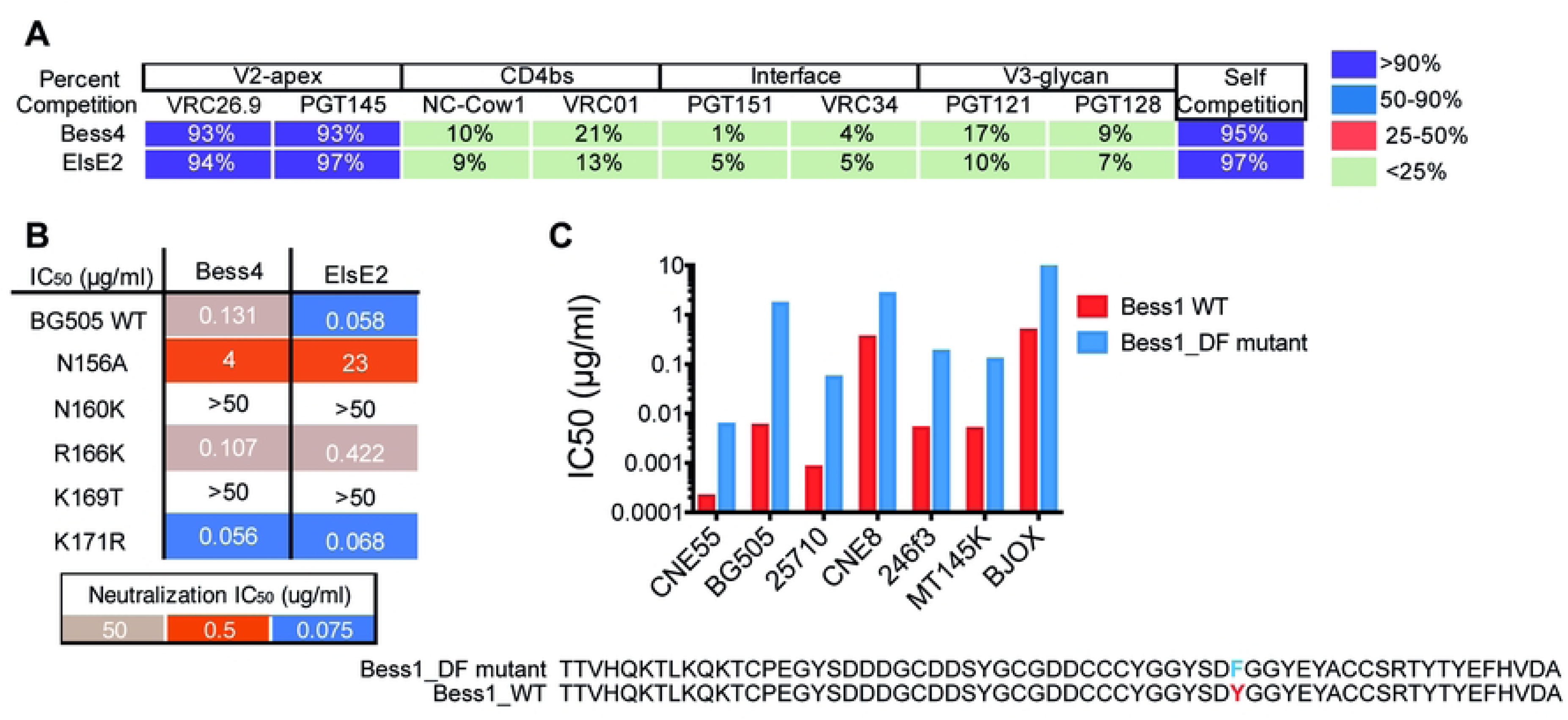
Cryo-EM structure of Bess4 in complex with BG505 SOSIP. **(A)** Structural similarities of the Bess7 and PGT145 CDR H3 regions. The Loop region of the Bess7 knob folds into a 14 residues β-hairpin, with i-i+3 residues DEYA at the distal tip. The long CDR H3 of human anti-HIV Fab PGT145 has a similar type I β-turn at its tip, with i-i+3 residues NETysG. **(B)** 3.3 Å cryo-EM reconstruction of Bess4 Fab in complex with BG505 SOSIP and RM20A3 Fab (used to increase orientation sampling). **(C)** Atomic model of Bess4 CDRH3 backbone with 3 pairs of disulfide bonds highlighted. **(D)** View of CDRH3 interaction with BG505 SOSIP as viewed down the trimer apex 3-fold axis. The three gp120 protomer chains are labeled A, B and C. **(E)** Comparison of Bess4 CDRH3 with human bnAbs PG9, PG16, CAP256-VRC26.25 and PGT145. Viewing plane is perpendicular to the viral membrane, with trimer apex at the top of the Fig. **(F)** Comparison of contact residues of BG505 SOSIP as defined by a buried surface area >10 Å^2^ between antibody CDRH3 and gp120. Residues with an asterisk denote contacts with N-linked glycan sugar(s).

To better understand and define the epitope recognized, we then chose Bess4 and ElsE2 as representatives to look at the effect of V2-apex epitope mutations on BG505 virus neutralization. Bess4 and ElsE2 neutralization titers were knocked out or reduced 30- to almost 400-fold for Bess4 and ElsE2, respectively, when mutations at position N156, N160, and K169 were introduced (Fig 6B). Human bnAbs PGT145, PG9, and CAP256-VRC26.9 are similarly affected by mutations at these residues on the V2-apex epitope, suggesting Bess and ElsE bind to the V2-apex of the trimer spike.

Finally, we wanted to determine whether tyrosine sulfation was present and necessary for neutralization in those antibodies. We used BLI to determine if all of the antibodies would bind to a mouse anti-sulfotyrosine antibody. The results were mixed. All of the Bess antibodies, except Bess4 and Bess6, had tyrosine sulfation detected, while none of the ElsE antibodies had detectable tyrosine sulfation (S3B Fig.). To investigate the specific amino acids responsible for the tyrosine sulfation, we mutated a DY motif in the wildtype Bess1 antibody to DF to eliminate sulfation (Fig 6C), which was confirmed by ESI mass spectrometry (S3C Fig.). The Bess1_DF mutant was tested for neutralization against seven viruses and compared to the Bess1 WT antibody. For almost every virus, neutralization was reduced but not eliminated (Fig 6C). Of further note, the CDRH3 sequences of Bess4 and Bess5 are very similar, but the more potent Bess4 does not have detectable tyrosine sulfation while Bess5 does, which could be due to a YD motif present in Bess5 but not Bess4. Overall, Bess and ElsE antibodies demonstrate a resemblance to other V2-apex targeting bnAbs in exclusively interactions with the trimer spike. However, they exhibit a varying dependence on tyrosine sulfation for virus neutralization.

### Structures of Bess Fabs reveal similarities in structure and binding to HIV trimers to human V2-apex bnAbs

Crystal structures were determined for 10 unliganded bovine Fab fragments, including ElsE1, ElsE2, ElsE5, ElsE6, ElsE7, ElsE8, ElsE9, ElsE11, Bess4 and Bess7 (S5 Fig., S4 Table). The ElsE Fabs all have CDRH3s with 54 amino acids and 4 cysteines, with a 1-3, 2-4 connectivity, Bess7 has a 61-amino acid CDRH3 with 8 cysteines (1-6, 2-5, 3-7, 4-8 connectivity), while the 55-amino acid CDRH3 knob region for Bess 4 (6 Cys, 1-3, 2-5, 4-6 connectivity) has no ordered electron density in the crystal structure. Flexibility in the stalk is evident in a comparison of the ElsE family of structures (S5 Fig.) that have similar CDRH3 knob sequences and structures, but very different orientations of knob domains with respect to the body of the Fab. For example, there are two Fabs in the ElsE11 asymmetric unit with a relative difference in the knob orientation of ∼114° (S5K and S5L Fig.). The ability of the knobs to twist and rotate with respect to the body of the Fab likely enhances the ability of bovine Fabs to access recessed epitopes. There is no convincing electron density to indicate tyrosine sulfation modifications in any of the antibody CDRH3 regions; however, BLI experiments indicate that Bess7, with 3 tyrosines in CDRH3, contains one or more sulfated tyrosines (S3 Fig.). Low sulfate occupancy or flexibility in the CDRH3 knob region may explain the lack of electron density for these groups. Ultralong bovine CDRH3 knob regions fold with 3 short β-strands connected by 2 loops (Loop1, Loop2) of varying lengths that can adopt a multitude of different secondary structures, including loops, helices and β-turns[22, 41]. The Loop2 region of the Bess7 knob forms a long, 14-residue β-hairpin, with residues DEYA forming a slightly distorted type I β-turn at the penultimate tip. This hairpin turn bears some resemblance to the tip of the long β-hairpin CDRH3 found in human HIV-1 bnAb PGT145, where residues NETysG form a type I β-turn at the tip (S5M Fig.). Bess7 is also notable for its interesting disulfide connectivity pattern between the 8 cysteines in its knob, with three (D gene positions 23, 24, 25) and two (D gene positions 39,40) sequential cysteines in the sequence (Fig 5A). Additionally, we screened Bess1-Bess4 Fabs for binding to BG505 SOSIP using negative stain EM. We found all of these Bess Fabs bound the V2-apex with a high degree of overlap, suggesting that despite differences in sequence, especially between Bess1-3 and Bess4, their mechanism of binding and approach angle to the SOSIP are similar (S5N Fig.).

To compare the binding interactions of the Bess antibodies with structurally characterized V2-apex human bnAbs, we determined a ∼3.3 Å cryo-EM structure of Bess4 Fab in complex with BG505 SOSIP, in which 41/57 CDRH3 residues were resolved, allowing for atomic modeling (Fig 7A, S6A-E Figs., S2 Table). The Bess4 CDRH3 forms a knob like structure, stabilized by 3 sets of disulfide bonds, and inserts down the Env apex 3-fold axis at an angle between three N160 glycans (Fig 7B, Fig 7C). This angled approach is similar to human bnAbs PG9, PG16 and CAP256-VRC26.25, in contrast to PGT145 which binds more perpendicularly with respect to the viral membrane (Fig 7D, S6F Fig.). Strikingly, despite coming from a different species and containing unique genetic properties relative to human bnAbs, Bess4 makes contacts with several residues also found at the interface with PG9, PG16, CAP256-VRC26.25 and PGT145 (Fig 7E, S6G Fig.). These structural similarities likely contribute to the broad neutralization measured for Bess4 and related antibodies in this study. 3D variability analysis of the EM data reveals significant movement of the rest of the Fab relative to the CDRH3, suggesting that CDRH3 makes all of the critical contacts with Env, and contributions from other parts of the antibody are minor (S6H Fig.,S1 Movie).

**Fig 7:**
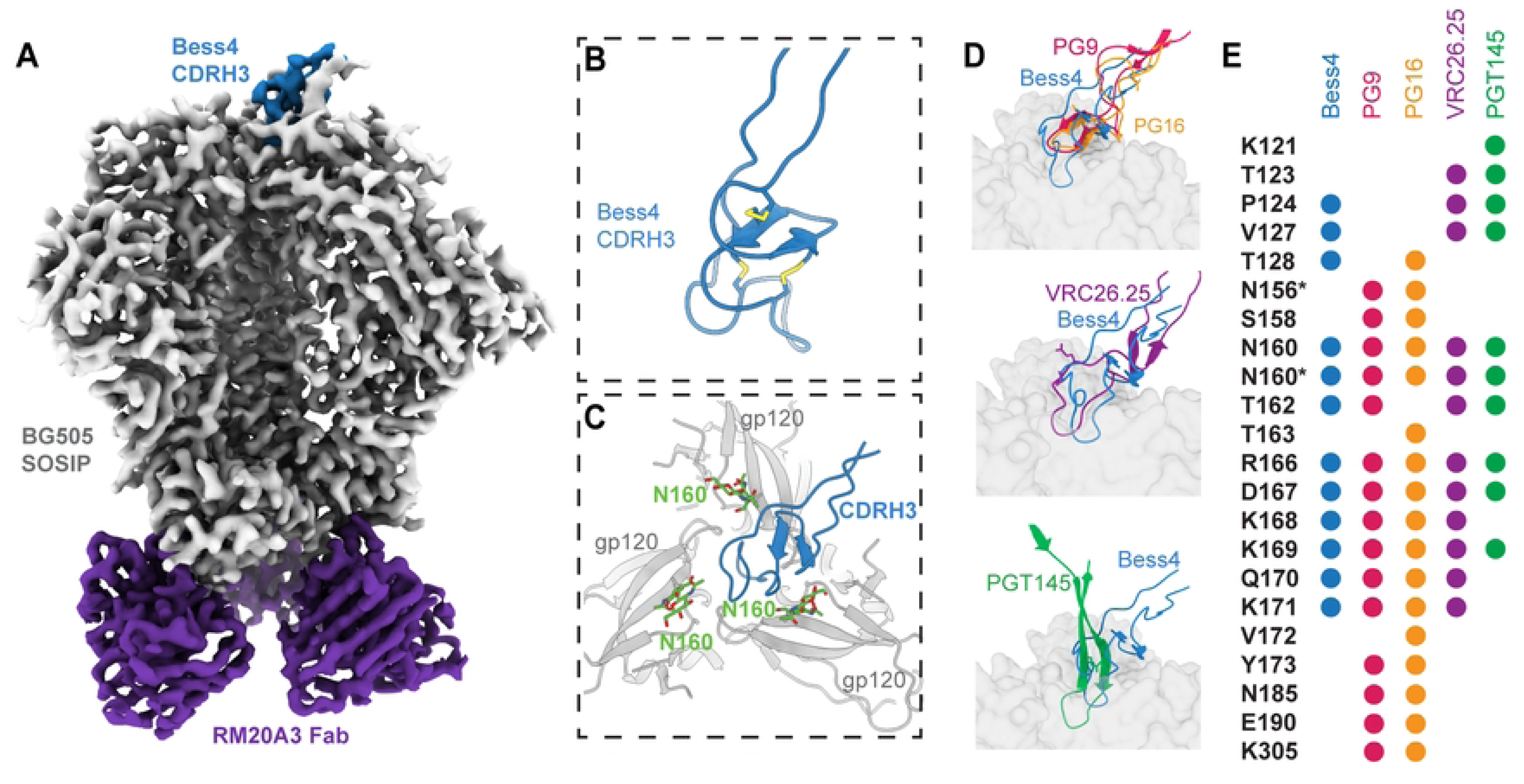

## Discussion

Previous studies have demonstrated that cows immunized with HIV Env immunogens can rapidly develop potent bnAbs to the CD4bs[47, 48]. This situation contrasts with other animal models where a sequential immunization regime involving different immunogens is likely required to initiate and then shepherd an antibody response toward bnAbs. An interesting question was whether the CD4bs had some specific features that were particularly amenable to ready recognition by cow antibodies with ultralong CDRH3s or whether the cow immune repertoire was inherently better suited to generate bnAbs to HIV. The study here, by describing cow bnAbs to the V2-apex site, suggests that there is an inherent ability of cow antibodies that allows for the development of broad neutralization of HIV, and this is related to the ultralong CDRH3 component of the cow repertoire. However, the bnAb sites are not equally accessed by cow antibodies, at least for the immunogens deployed here. The V2-apex serum neutralizing titers induced were lower and required more immunizations than those induced to the CD4bs and furthermore, the V2-apex bnAbs described here are somewhat less potent and broad than for example the CD4bs antibody NC-Cow1.

Another rationale for this study was that the cow repertoire includes not only ultralong CDRH3s but also a higher frequency of long CDRH3s of a length similar to those of V2-apex bnAbs isolated from HIV-infected donors. It was therefore of interest to determine whether appropriate trimer immunization might generate bnAbs from the long CDRH3 repertoire of the cow. In fact, all the bnAbs that we isolated were from the ultralong CDRH3 compartment of the cow repertoire. This observation suggests that the ultralong CDRH3 antibodies have an advantage over the long CDRH3 antibodies in recognizing HIV Env and are more competitive in the selection process involved in the initiation and maturation of antibodies that lead to a bnAb response. One caveat is that long and short CDRH3 antibodies in aggregate may have been placed at a disadvantage in our study, as it is unclear if they can always pair with the universal cow light chain although most were tested with their native light chain.

A further rationale for this study was that, if it was possible to produce cow antibodies directed to bnAb sites other than the CD4bs, then the antibodies might be useful in cocktails to generate reagents with a greater activity across the diversity of HIV and to help restrict neutralization escape. In particular, the knob structures associated with ultralong CDRH3 antibodies have intrinsic potent bnAb activity and, therefore, represent small units that could be incorporated into different frameworks. For example, an effective microbicide using bnAbs or their derivatives would require a cocktail and the maintenance of potency and stability at low pH. Previous studies showed that the ultralong CDRH3 cow bnAb, NC-Cow1, was able to maintain gp120 binding in simulated vaginal fluid at pH 4.5[47]. The ultralong knob domain itself has been isolated from the full IgG framework and shown to be able to bind to antigens of interest with similar affinity to the whole IgG[59, 60]. Other studies have engineered knobs into protein loops to produce novel multivalent molecules[61-65]. In addition to binding, a recent study demonstrated that recombinant knob domains alone are able to potently neutralize SARS-CoV-2^[34]^. Therefore, a multivalent construct based on cow bnAb CDRH3 knobs, or knob cocktails, may be worthy of investigation.

In summary, immunization of animal models and humans with native-like Env trimers does not generate bnAbs as detected in notable serum titers with the single exception of immunization of cows. This exception arises because of the propensity of ultralong CDRH3 cow antibodies to penetrate the glycan shield and specifically recognize the conserved regions of two bnAb sites on HIV Env. The knob structures responsible for broad neutralization form protein units of exceptional stability that may find therapeutic applications in appropriate formats.

## Materials and Methods

### Lead contact

Further information and requests for resources and reagents should be directed to and will be fulfilled by the lead contact, Devin Sok.

### Materials availability

All unique reagents in this study are available from the lead contact with a completed Materials Transfer Agreement.

### Data and code availability

Fab crystal structure data was deposited into the Protein Data Bank using the following codes: ElsE1 (PDB: 8V4I), ElsE2 (PDB: 8VBJ), ElsE5 (PDB: 8VBK), ElsE6 (PDB: 8VBL), ElsE7 (PDB: 8VBM), ElsE8 (PDB: 8VBN), ElsE9 (PDB: 8VBO), ElsE11 (PDB: 8VBR), Bess4 (PDB: 8VBP), Bess7 (PDB: 8VBW). The structure of Bess4 in complex with BG505 SOSIP was deposited into the Electron Microscopy Data Bank under accession code EMD-41498 and the Protein Data Bank under accession code PDB: 8TQ1. Additional data related to this paper is available from the lead contact upon request. This paper does not report original code.

## EXPERIMENTAL MODEL AND SUBJECT DETAILS

### Cow Model: *Bos taurus*

#### Cell Lines

Cells were HEK-derived and included 293T, 293F, and Expi293 cells. TZM-bl cells are a He-La derived cell line that express CD4 receptor and CXCR4 and CCR5 chemokine co-receptors and luciferase and β-galactosidase genes under the control of the HIV-1 promoter. Both 293T and TZM-bl cell lines were maintained and used in complete Dulbecco’s Modified Eagle Medium (complete DMEM) which is comprised of high-glucose Dulbecco’s Modified Eagle Medium, 2 mM L-glutamine, 10% fetal bovine serum, and 1x Penicillin-Streptomycin. The 293T and TZM-Bl cell lines were maintained at 37°C and 5% CO_2_. 293F and Expi293 cells were maintained in Gibco Freestyle 293 Expression Medium (Thermo Fisher) and Gibco Expi293 Expression Medium (Thermo Fisher) at 37°C and 10% CO_2_ with shaking at 120 RPM.

### Method details

#### Cow immunizations

Antigen-specific B cell responses were primed and expanded by immunization of *Bos taurus* calves. The calves were bled from the jugular vein as often as once a week from the beginning of the immunizations and bi/tri-monthly during the last 3 months of the immunizations. Blood for sera and for isolation of PBMCs were collected at the same time and terminal bleeds and splenocytes were collected at study termination. The calves and samples were not randomized or blinded. The calves were immunized with the appropriate immunogen at a dose of 200 µg per calf (formulated in Montanide ISA 201 adjuvant: Seppic, France), for all priming and boosting steps except the final boost. The immunogens were inoculated intradermally (200µl per site) on both sides of the neck region. The final boost was done with a SOSIP cocktail containing BG505.664, CRF250, WITO, and ZmZm containing 100 µg of each SOSIP formulated as above. Group 1 with cow-485 and cow-16157 were immunized with MT145K SOSIP.664 on days 0, 37, and 79 then with C108 SOSIP.664 on days 142 and 226. The final immunization on day 336 was done with a SOSIP.664 cocktail consisting of BG505, CRF250, WITO, and ZmZm. Group 2 with cow-488 and cow-491 were immunized with C108 SOSIP.664 on days 0, 37, 79, 142, and 226. The final immunization on day 336 was done with a SOSIP.664 cocktail consisting of BG505, CRF250, WITO, and ZmZm.

#### Sera IgG purifications

Sera were heat-inactivated in a 56°C water bath for 30 minutes and spun at maximum speed in a table-top centrifuge for 20 minutes. Sera were subsequently incubated with an equal volume of dry Protein G Sepharose beads (GE) and twice the volume of PBS. This was then incubated overnight at 4°C on a nutator. The next day the beads were washed with 10x the volume of PBS, eluted with IgG elution buffer (Pierce), and neutralized with 2M Tris pH 9.0. The eluted fraction was buffer exchanged into PBS and concentrated into the original sera volume. Samples were filtered through a 0.45 µm filter.

#### Pseudovirus neutralization assays and pseudovirus production

Replication incompetent HIV pseudovirus was produced in HEK293T cells. They were co-transfected with plasmids containing HIV Env and PSG3ΔEnv backbone using a 1:2 ratio. FuGENE (Promega) was used as a transfection reagent in Opti-MEM (Thermo Fisher) or Transfectagro (Corning). Supernatants from cell cultures were harvested 48-72 hr post transfection and sterile filtered with a 0.22 mM filter. Pseudovirus was either used neat or concentrated as needed using 50k amicon concentrators. Pseudovirus was either frozen or used fresh. Frozen pseudovirus was titrated to determine concentration needed for use. Neutralization assays were performed by incubating 25 µl of monoclonal antibodies or IgG purified from sera with 25 µl of pseudovirus for 1 hr at 37°C in full-area 96-well tissue culture polystyrene microplates (Corning). After 1 hr 20 µl of TZM-bl target cells were added at a concentration of 0.5 million cells/mL with 40 µg/mL of dextran final[66]. Cells were grown in a humidified incubator for 24 hr, then 130 µl of media was added to all wells. 48-72 hr after cells were added, plates were read on a luminometer (Biotek) by adding lysis buffer combined with Bright-Glo (Promega). Neutralization was measured in duplicate wells and pseudovirus and cell controls were averaged for analysis. Neutralization of purified IgG from serum samples was tested starting at 1:35 dilutions followed by 3-fold serial dilutions. Monoclonal antibody neutralization assays were diluted starting at 5-100 µg/mL with a 4 to 10-fold serial dilution depending on what was appropriate for the assay. All neutralization titers are reported as ID_50_ (1/dilution) or IC_50_ (µg/mL) titers. nAb titer data panels are shown as geometric mean titers. ZM233 was used in place of ZMZM for neutralization assays. Analysis and graphing were done in Prism software.

#### SOSIP trimer purification

MT145K, C108, CRF250, BG505.664dHis, WITO, 25710, CRF250, and ZM233 SOSIP were expressed in HEK293F cells and purified as described previously described[49, 67]. Briefly, SOSIP was transfected with 300 µg of trimer DNA, 150 µg of Furin DNA, and 1.5 mL of PEI MAX 40000 (Polysciences) in Transfectagro. Five to six days after transfection supernatants were harvested and purified using a G*alanthus nivalis* lectin (Vector Labs) column or PGT145 affinity columns which are made by coupling CnBr-activated Sepharose 4B beads (GE Healthcare) to PGT145 bnAb antibody as described previously[67]. Purified proteins were further purified with size exclusion chromatography columns Superdex 200 10/300 GL column (Cytvia) in TBS or PBS. Trimers were frozen for storage at -80°C and then thawed and tested with ELISA or BLI for antigenicity with a range of known non-neutralizing and neutralizing HIV specific antibodies. SOSIP used as sorting baits were biotinylated using the Pierce Biotinylation Kit using the manufacturer’s protocol and tested for biotinylation via ELISA.

#### Single cell sorting

Cow PBMCs and splenocytes were sorted as previously described with study specific modifications[53]. Cow cells were thawed in a 37°C water bath until only a small pellet of ice remained. Cells were resuspended with 10 mL of pre-warmed buffer consisting of 50% RPMI (Gibco) and 50% FBS. Cells were washed by centrifuging for 5 min at 400g. Supernatant was discarded and the cell pellet was gently resuspended in 5 mL of cold FACS Buffer (1x DPBS with 1% FBS, 1mM EDTA, and 25 mM HEPES). Cells were counted and washed, the FACS Buffer was then removed. FACS antibody mastermix was added per 10 million cells. FACS antibody master mix per 10 million cells consisted of 3.75 µg of FITC labeled goat-anti cow antibodies (Abcam) and SOSIP baits diluted in FACS Buffer. Baits for the first three sorts used 55.5 pM of each biotinylated and stained SOSIP which was coupled with fluorophore conjugated streptavidin in a 4:1 molar ratio for 1 hr on ice in the dark. The fourth sort used 200 nM of biotinylated SOSIP coupled in a 1:2 ratio with a streptavidin fluorophore. Additionally, one of these SOSIPS was combined with PGT145 Fab after staining to form a SOSIP-Fab complex. The cell and master mix combination was incubated on ice in the dark for 30 minutes. After 30 minutes 1 mL of 1:300 diluted FVS510 Live/Dead stain (BD Biosciences) was added per 10 million cells and incubated in the dark for 15 min. Cells were washed in 10 mL of FACS buffer and resuspended in 500 µl per 10 million cells of FACS buffer. The cells were filtered into a 5 mL round bottom FACS tube with a cell strainer cap (Falcon). The fourth sort used MT145K SOSIP coupled with streptavidin-PE (Invitrogen) and MT145K SOSIP complexed with Fab and coupled to streptavidin AF-647 (Invitrogen). The first and second sort selected for cells that had double positive binding to two of the three SOSIP baits which were 25710 (BV421, BD Biosciences), CRF250 (PE, Invitrogen), and MT145K (APC, Invitrogen). The third sort selected for cells that demonstrated double positive binding for the same SOSIP (MT145K or 25710) coupled individually to PE (Invitrogen) or AF-647 (Invitrogen). Cells from the first three sorts were sorted into 96-well plates containing 20 µl/well of lysis buffer on FACS ARIA III BD FACS sorter and immediately frozen on dry ice. Lysis buffer consists of 2.5 mM RNaseOUT (Invitrogen), 1.25x SuperScript IV Reverse Transcriptase Buffer (Invitrogen), 6.25 mM DTT, and 1% Igepal CA-630 (Sigma-Aldrich). The fourth sort selected cells positive to binding only MT145K SOSIP (PE) and did not have affinity for MT145K-PGT145 Fab Complex (AF-647). Cells were sorted into dry 96-well plates that were immediately frozen on dry ice. Data was analyzed using FlowJo.

#### Single cell PCR amplification and cloning

cDNA was generated from single cells sorted into plates using RT-PCR. RT-PCRs were done two ways depending on if the plate was dry or had lysis buffer. The plates with lysis buffer had 9 µl of RT-PCR buffer added to them consisting of 100 U of SuperScript IV Reverse Transcriptase Enzyme (Invitrogen), 1.1 mM dNTPs(New England Biolabs), 0.44x SSIV buffer(Invitrogen), and 0.375 µg of random hexamers (Gene Link). Each well in the dry plates received 15 µl of 3.33 mM RNaseOut (Invitrogen), 1x SuperScript IV Reverse Transcriptase Buffer (Invitrogen), 100 U of SuperScript IV Reverse Transcriptase Enzyme (Invitrogen), 0.225 µg of random hexamers (Gene Link), 0.33 mM dNTPs (New England Biolabs), 4.3 mM DTT, and the rest supplemented by RNAse Free Water.. RT PCR used the following PCR program: 10min at 42°C, 10 min at 25°C, 10 min at 50°C, followed by 5 mins at 94°C. Nested PCRs were done to amplify heavy chain and lambda light chain variable regions using multiple primers. PCR1 used primers CowVHfwd1: CCCTCCTCTTTGTGCTSTCAGCCC, CowIgGrev1: GTCACCATGCTGCTGAGAGA, and CowIgGrev2: CTTTCGGGGCTGTGGTGGAGGC. PCR1s were done in 20 µl reactions per well/single cell using 3 µl of RT PCR product, 1x HotStarTaq Mastermix (New England Biolabs), 0.2 µM of forward primers total, and 0.2 µM of reverse primers total, with the rest supplemented by RNAse Free Water. The following PCR program was used: 15s at 95°C, 49 cycles of 30s at 94°C, 30s at 55°C, and 60s at 72°C followed by an extension for 10 min at 72°C. PCR2 was done using heavy chain primers CowVHPCR2:CATCCTTTTTCTAGTAGCAACTGCAACCGGTGTACATTCCMAGGTGCAGCTGCRGGAGTC and CowIgGRevPCR2: GGAAGACCGATGGGCCCTTGGTCGACGCTGAGGAGACGGTGACCAGGAGTCCTTGGCC. Light chains were amplified with primers L Leader 2 F: CACCATGGCCTGGTCCCCTCTG, L Leader 15F: GGAACCTTTCCTGCAGCTC, L Leader 16 F: GCTTGCTTATGGCTCAGGTC, L Leader 35F: GACCCCAGACTCACCATCTC, L Leader 45 F: AGGGCTGCGGGCTCAGAAGGCAGC, L Leader 55 F: CTGCCCCTCCTCACTCTCTGC, and Cow_LC_rev1: AAGTCGCTGATGAGACACACC. PCR2 was done for the light chain using primers Fwd-PCR2-LC: CATCCTTTTTCTAGTAGCAACTGCAACCGGTGTACACCAGGCTGTGCTGACTCAG and LC_REV_const-PCR2: GTTGGCTTGAAGCTCCTCACTCGAGGGYGGGAACAGAGTG. The PCR 2 was used to add tags for Gibson assembly. PCR2 introduced homology to the cut ends of the variable regions for later cloning and were done in 25 µl reactions per well/single cell using 2 µl of PCR product from PCR1, 0.5 U of Phusion Enzyme (Thermo Fisher), 0.2 mM dNTPs at 10 mM each (New England Biolabs), 1 µM of forward primers total, 1 µM of reverse primers total,1.5 mM MgCl_2_, 1x HF Buffer (Thermo Fisher), and the rest supplemented with RNAse Free Water using the following PCR program: 30s at 98°C then 34 cycles of 10s at 98°C, 40s at 72°C, followed by an extension for 5 min at 72°C. PCR products were resolved with E-gel 96 Agarose Gels 2% (Invitrogen) to determine if PCR chains were properly amplified. PCR products were purified using SPRIselect beads (Beckman Coulter) and amplified variable regions were sanger sequenced and results were analyzed using the International ImMunoGeneTics (IMGT) Information System (www.imgt.org) V-quest[68]. All isolated variable regions from PCR products were constructed into human antibody expression vectors with appropriate IgG1 or Ig lambda constant domains using high throughput NEBuilder HiFi DNA assembly system. 10 ng of restriction enzyme digested backbone was combined with 20 ng of SPRI cleaned PCR product followed by the appropriate volume of 2x HiFi master mix (New England Biolabs). Gibson assembly reactions were transformed into DH5 Alpha Competent Cells (Biopioneer) and single cultures were grown up and purified using QIAprep Spin Miniprep Kit (Qiagen).

#### Antibody and Fab production and purification

Monoclonal antibodies were transiently expressed in Expi293 and 293F cells. Monoclonal cow antibodies were expressed with bovine V_L_, V_H_, and human C_L_, C_H_ regions. In Expi293 cells heavy chain and light chain pairs were co-transfected at a ratio of 1:2.5 with FectoPRO (Polyplus) in Opti-MEM (Thermo Fisher) or Transfectagro (Corning). After 22-24 hr cultures were supplemented with 300 mM of sodium valproic acid solution (Sigma-Aldrich) and 45% D-(+)- glucose solution (Sigma-Aldrich). In 293F cells, heavy chain and light chain DNA were added in a 1:1 ratio with PEI as a transfection reagent in Opti-MEM (Thermo Fisher) or Transfectagro (Corning). The supernatant was collected 4-5 days after transfection and sterile filtered through an 0.22 or 0.45 mm filter. Sterile supernatants were used for screening or whole IgGs were purified using Protein A Sepharose (GE Healthcare) or Praesto AP (Purolite).

Fabs were produced either with papain digestion or recombinantly. For digestion from monoclonal antibodies, Fabs were prepared with the Pierce Fab Preparation Kit (Thermo Fisher) following the manufacturer’s protocol for human Fab digest. Fab from polyclonal IgG purified from sera were prepared using Pierce Fab Preparation Kit with each digest at 0.5 mg/mL and a digest time of 16 hr.

Fabs with bovine V_L_, V_H_, and human C_L_, C_H_1 regions were expressed by transient transfection in Expi293F cells (Thermo Fisher) following the manufacturer’s standard protocol. Fabs were purified from the media using CaptureSelect CH1-XL resin (Thermo Fisher) and further purified by size exclusion chromatography on a Superdex 200 16/600 column (Cytiva) with running buffer 20mM Tris, 150mM NaCl, pH 8.

#### High throughput antibody screening

Antibodies were transiently expressed in 3 mL cell cultures of Expi293 cells using 24 deep-well plates (Thermo Fisher). After 4 days, supernatants were harvested and filtered with 0.22 or 0.45 µm membrane filters and were used neat for initial screening of expression, SOSIP binding, and neutralization. We used an antibody detection ELISA to determine if antibodies were expressed and able to bind to trimer (see ELISA section). If antibodies were expressed, neat supernatant was tested for neutralization. Neutralization assays were done as previously described using autologous CRF250 and C108 viruses and 12-virus panel virus CNE55. If any neutralization was detected, antibodies were expressed in 30 to 50 mL cultures and purified using Protein A Sepharose (GE Healthcare) or Praesto AP (Purolite). Purified monoclonal antibodies were tested for neutralization and subsequent analysis. IgHV1-7 derived heavy chains were paired with universal V30 light chains and native light chains if they were recovered. Others were only screened with native light chain when recovered[22, 41]. The neutralization was not different between purified monoclonals tested with native and universal light chain and all large-scale production of Bess monoclonals were expressed with V30 light chains. ElsE antibodies were all expressed and further tested with their native light chains.

#### BioLayer interferometry (BLI) assays

BLI assays were performed using polypropylene black 96-well microplate (Greiner) on an Octet RED384 instrument at 30°C, in the octet buffer comprising PBS with 0.05% Tween. All reagents were diluted with octet buffer. Antibody competition assays were done in-tandem to determine the binding epitopes of isolated monoclonal antibodies. 222 nM of biotinylated 25710 SOSIP was captured with Streptavidin (SA) biosensors (Sartorius) for 10 min and transferred to Octet buffer for 1 min to wash off unbound SOSIP. Sensors were then moved into monoclonal Bess and ElsE antibodies at a concentration of 400 nM for 10 min and washed off in octet buffer for 1 min. Biosensors were then moved into known antibodies at a concentration of 200 nM for 5 min. The percent inhibition in binding was calculated with the formula: Percent binding inhibition (%) = 1- (competitor antibody binding response in presence of saturating antibody) / (binding response of the competitor antibody without saturating antibody). BLI assays to determine binding of Bess and ElsE to anti-sulfotyrosine antibodies were performed using polypropylene black 384-well microplate (Greiner) on an Octet RED384 instrument at 30°C in octet buffer. We used anti-human IgG Fc Capture (AHC) biosensors (Sartorius) to capture monoclonal antibodies at a concentration of 50 µg/mL for 5 min and washed with octet buffer. After antibodies were captured, the biosensor was moved into 1:50 diluted mouse Anti-Sulfotyrosine Antibody, Clone Sulfo-1C-A2 (MilliporeSigma) to detect binding for 200 sec. All dilution and incubation steps were performed in 1x PBS with 0.05% Tween. Analysis and graphing were done in Prism software.

#### ELISA assays

All ELISA assays were done with half-area 96-well high binding ELISA plates (Corning). All washes were done 3x with PBS containing 0.05% Tween20 and all antibodies, except those used for coating, were diluted in PBS containing 1% BSA and 0.025% Tween20. All reagents added to the plate were in 50 µl volumes per well, except blocking which was done in 150 µl volumes. All steps and incubations were done at room temperature unless otherwise stated. All plates were developed with phosphatase substrate (Sigma-Aldrich) diluted in alkaline phosphatase staining buffer (pH 9.8), according to manufacturer’s instructions. Optical density at 405 nm was read on a microplate reader (Molecular Devices). Sera competition ELISA assay plates were coated with PBS containing 250 ng of anti-6x His tag monoclonal antibody (Invitrogen) and left covered overnight at 4°C. The next day plates were washed and blocked with 3% BSA for 1-2 hr at 37°C. After washing 125 µg of his-tagged BG505 was added to all wells for 1 hr. IgG purified from sera were added using serial dilutions starting at 1:20 with seven 4-fold dilutions and incubated for 1 hr. Biotinylated monoclonal antibodies with known epitopes were then added at a constant concentration, depending on their previously measured EC_70_ value, for 1 hr. After washing, alkaline phosphatase conjugated to streptavidin at a dilution of 1:1000 (Jackson ImmunoResearch) was added to plates and allowed to incubated for 1 hr. Plates were washed, substrate was added, and plates were read as mentioned previously. Biotinylated antibody EC_70_’s were measured using the same methods, but without the addition of purified IgG from sera. ELISA assays for gp120 binding were done by coating each plate with PBS containing 250 ng of BG505 gp120 and left covered overnight at 4°C. The next day plates were washed and blocked with 3% BSA for 1-2 hrs. Monoclonal antibodies were added using serial dilutions starting at 20 µg/mL with seven 6-fold dilutions and incubated for 1 hr. After washing, alkaline phosphatase conjugated anti-human F(ab’)_2_ secondary antibodies (Jackson ImmunoResearch) at a dilution of 1:1000 were added to plates and incubated for 1 hr. After a final wash, substrate was added, and plates were read as mentioned previously. High throughput antibody expression assays used ELISA for the detection of antibody expression and BG505 SOSIP binding in harvested expi293 transfected supernatant. Plates were coated overnight at 4°C with PBS containing 100 ng of AffiniPure Fragment Goat Anti-Human IgG, F(ab’)₂ (Jackson ImmunoResearch). Next day the plates were washed and blocked with 3% BSA for 2 hr at 37°C. 50 µl of neat supernatant were added to each well for 1 hr. After washing Alkaline phosphatase conjugated anti-human Fc secondary antibodies (Jackson ImmunoResearch) was added at a dilution of 1:1000 for 1 hr. After a final wash, substrate was added, and plates were read as mentioned previously. The trimer binding ELISA assay used plates coated with 100 ng of 6x His tag monoclonal antibodies (Invitrogen) in PBS and left overnight at 4°C. The next day plates were washed then blocked with 3% BSA for 2 hr at 37°C. Plates were washed, then 125 µg of 6x His-tagged BG505 SOSIP were incubated in each well for 1 hr. Plates were washed and neat supernatant from high throughput antibody transfections were added for 1 hr. Plates were washed and alkaline phosphatase conjugated anti-human F(ab’)_2_ secondary antibodies (Jackson ImmunoResearch) at a dilution of 1:1000 were added to each well. After a final wash, substrate was added, and plates were read as mentioned previously. All data and graphs were analyzed using Prism software.

#### Phylogenetic tree building

Phylogenetic trees were created using Geneious Prime software. The full variable region or the CDRH3 of NC-Cow1, Bess, and ElsE amino acids starting were aligned with Clustal Omega 1.2.3. The alignment was made into a tree using Geneious Tree Builder software. We used Jukes-Cantor for the genetic distance model and the Neighbor-Joining Tree Build Method.

#### X-ray crystallography

Recombinant Fabs were screened for crystallization with our robotic Rigaku CrystalMation system and JCSG 1-4 crystal screens at 4° and 20°. Crystallization conditions for crystals used for data collection are included in S1 Table. Data were collected at synchrotron beamlines at SSRL, ALS, and APS (S4 Table) and processed with HKL-2000^[69]^. Structures were determined by molecular replacement using model coordinates from Fab BLV1H12 (PDB 4K3D) and Phaser[22],[70]. Models were adjusted and rebuilt in Coot and refined with Phenix.refine[71, 72]. Final data collection and refinement parameters are listed in S5 Table.

#### Cryo-electron microscopy

0.2 mg of BG505 SOSIP was incubated with 0.15 mg of Bess4 Fab (from this study) and 0.3 mg of base-directed RM20A3 Fab (to increase angular sampling in cryo-EM) overnight at room temperature and purified the following morning using a HiLoad 16/600 Superdex 200 pg (Cytiva) gel filtration column[73]. The complex was then concentrated to 5 mg/mL for application onto cryoEM grids. Cryo grids were prepared using a Vitrobot Mark IV (Thermo Fisher Scientific). The temperature was set to 4°C and humidity was maintained at 100% during the freezing process. The blotting force was set to 1 and wait time was set to 10 s. Blotting time was varied from 3 to 4 s. Detergents lauryl maltose neopentyl glycol (LMNG; Anatrace) or n-Dodecyl-β-D-Maltoside (DDM; Anatrace) at final concentrations of 0.005 or 0.06 mM, respectively, were used for freezing. Quantifoil R 1.2/1.3 (Cu, 300-mesh; Quantifoil Micro Tools GmbH) or UltrAuFoil 1.2/1.3 (Au, 300-mesh; Quantifoil Micro Tools GmbH) grids were used and treated with Ar/O_2_ plasma (Solarus plasma cleaner, Gatan) for 8 sec before sample application. 0.5 µL of detergent was mixed with 3.5 µL of samples and 3 µL of the mixture was immediately loaded onto the grid. Following blotting, the grids were plunge-frozen into liquid nitrogen-cooled liquid ethane. Samples were loaded into a Thermo Fisher Scientific Titan Krios operating at 300 kV. Exposure magnification was set to 130,000x with a pixel size at the specimen plane of 1.045 Å. Leginon software was used for automated data collection[74]. Micrograph movie frames were motion corrected and dose weighted using MotionCor2 and imported into cryoSPARC for the remainder of data processing. CTF correction was performed using cryoSPARC Patch CTF[75, 76]. Particle picking was performed using blob picker initially followed by template picker. Multiple rounds of 2D classification and 3D ab-initio reconstruction were performed prior to 3D non-uniform refinement with global CTF refinement. Due to substoichiometric binding and high flexibility of the Bess4 antibody with respect to the trimer, particles were symmetry expanded (C3 symmetry), subjected to 3D Variability with a mask over the trimer apex, and 3D Variability clustering analysis. Clusters with visible Fab density were pooled, duplicate particles removed, and a final Non-Uniform Refinement was performed (C1 symmetry) with global resolution estimated by FSC 0.143. Final data collection and processing stats are summarized in S5 Table. The Bess4 Fab x-ray structure (this study) was docked into the EM map along with a model of BG505 SOSIP in complex with RM20A3 (PDB 6x9r) in UCSF Chimera[77]. The missing CDRH3 residues from the x-ray structure were built manually in Coot 0.9.8 and real space refinement using Rosetta and Phenix^[78-80]^. The final model includes 41 of the 55 CDRH3 residues of Bess4, and all other regions of the Fab were trimmed due to lower map resolution resulting from high flexibility. The final model was validated using MolProbity and EMRinger in the Phenix suite, and statistics are summarized in S5 Table. The map and model have been deposited to the Electron Microscopy Data Bank and Protein Data Bank, respectively with accession codes summarized in S5 Table.

#### Production of cDNA from PBMCs

Total RNA was extracted from PBMCs using RNeasy mini kit (Qiagen) from D359. Following RNA isolation, cDNA was made using Superscript IV reverse transcriptase (Thermo Fisher). For the cDNA reaction, 549 ng (3 µL) of RNA was used as template, along with 167 nM (1 µL) of JH Bov rvs primer (TGAGGAGACGGTGACCAGGAGTC), 1 µL dNTP, and 9 µL of DNase and RNase-free water. This mixture was incubated at 65°C for 5 min. 5X buffer was vortexed and spun down, after which 4 µL of 5X buffer was mixed with 1 µL DTT, 1 µL ribonuclease inhibitor, and 1 µL SuperScript IV. All 8 components were then mixed, followed by incubation at 52°C for 10 min and then 80°C for 10 min.

#### PCR and sequencing

PCR to amplify cow antibody heavy chains was set up as follows: 2.5 µL of 10 µM forward primer, 2.5 µL of 10 µM reverse primer, 2 µL of cDNA, 25 µL of 2X Q5 Hot-start High-fidelity Master Mix (NEB), 18 µL of DNase and RNase-free water. The cycling parameters were as follows: 98°C for 1 min, 98°C for 10 s, 72°C for 1 min, repeat Steps 2 and 3 34 more times, 72°C for 2 min, hold at 4°C. The CDR1F forward primer, which anneals to VH1-7, has the following sequence: TTGAGCGACAAGGCTGTAGGCTG. The VH4 forward primer, which anneals to all cow VH genes, has the following sequence: CTGGGTCCGCCAGGCTCC. The JH Bov rvs reverse primer (see above), which anneals to cow JH2-4, was used as the reverse primer in all reactions. Following PCR, the products were purified using the QIAquick PCR Purification Kit (Qiagen) according to the manufacturer’s protocol. PCR products were then sent to GeneWiz for Amplicon-EZ sequencing, which was performed on an Illumina platform to generate ∼100,000 2 X 250 bp paired-end reads per sample.

#### Computational analysis of sequences

Once sequences were obtained from GeneWiz, non-functional sequences were removed using SAS. Sequences were considered non-functional if they did not start with the conserved VRXA motif, where X represents any amino acid, or if they did not contain the conserved WG motif at the end of CDRH3. In addition, any sequence was considered non-functional if it contained a premature stop codon in the VH region or if it was less than 220 bp (73 amino acids) long. Using the R computer programming language, the abundance of each amino acid sequence was determined. Analysis of CDRH3 sequences and ultralong antibodies was also performed in R.

## Acknowledgements

Use of the Stanford Synchrotron Radiation Lightsource, SLAC National Accelerator Laboratory, is supported by the U.S. Department of Energy, Office of Science, Office of Basic Energy Sciences under Contract No. DE-AC02-76SF00515. The SSRL Structural Molecular Biology Program is supported by the DOE Office of Biological and Environmental Research, and by the National Institutes of Health, National Institute of General Medical Sciences (P30GM133894). The contents of this publication are solely the responsibility of the authors and do not necessarily represent the official views of NIGMS or NIH. GM/CA@APS has been funded by the National Cancer Institute (ACB-12002) and the National Institute of General Medical Sciences (AGM-12006, P30GM138396). This research used resources of the Advanced Photon Source, a U.S. Department of Energy (DOE) Office of Science User Facility operated for the DOE Office of Science by Argonne National Laboratory under Contract No. DE-AC02-06CH11357. The Eiger 16M detector at GM/CA-XSD was funded by NIH grant S10 OD012289. The Berkeley Center for Structural Biology is supported in part by the Howard Hughes Medical Institute. The Advanced Light Source is a Department of Energy Office of Science User Facility under Contract No. DE-AC02-05CH11231. The Pilatus detector on 5.0.1. was funded under NIH grant S10OD021832. This work was supported by National Institutes of Health grants R61 AI161818 (RA) and the National Institute of Allergy and Infectious Diseases (NIAID) Consortium for HIV/AIDS Vaccine Development (CHAVD; UM1AI144462) (DRB, DS, ABW and IAW).

## Supplemental fig, table, and movie titles and legends

**S1 Fig. Four single B cell sorts were used to isolate monoclonal antibodies of interest.** Each sort done on samples from both cow-488 and cow-485

**S2 Fig. Phylogenetic trees of both the full variable region and CDRH3 of ElsE and Bess antibodies.** Else, Bess, and NC-Cow1 variable regions (top) starting at FR1 and the CDRH3 (bottom) were aligned and categorized into a phylogenetic trees. Bess antibodies are colored in red, ElsE in blue, and NC-Cow1 in purple.

**S3 Fig. Bess and ElsE antibodies were evaluated for their ability to bind to monomer and whether they were tyrosine sulfated.**

**(A)** All Bess and ElsE antibodies were measured for binding to BG505 gp120 using ELISA. Antibody controls include positive controls VRC01 and F425, and negative control CAP256-VRC26.9 (VRC26.9). Bess1-8, ElsE1-11 mAbs, and CAP256-VRC26.9 showed no detectable binding to BG505 gp120. **(B)** Bess and ElsE series mAbs were evaluated for their ability to bind to a mouse derived anti-sulfotyrosine antibody using BLI. NC-Cow1 and Den3 were used as negative controls. CAP256-VRC26.9 and PGT145 were used as positive controls. Positive and negative control lines are shown as dashes. Table showing which antibodies bound are shown on the right. **(C)** To evaluate loss of tyrosine sulfation, ESI mass spectrometry was used to compare the mass difference of Bess1_WT and Bess1_DF Mutant. ESI mass spectrometry spectra are shown. Below is a chart with a comparison of the mass differences and expectations for the mAbs, amino acids, and tyrosine sulfation sizes sulfation are summarized at the bottom.

**S4 Fig. SPR Data of ElsE and Bess antibodies binding to BG505 SOSIP.**

Summarized results of BG505 binding to Bess and ElsE mAbs via Protein A capture, multi-cycle method. SPR experiments are shown in black and best global fits are shown in red. 1:1 Langmuir binding model and stead-state were used to calculate the association (ka) and dissociation (kd) rate constants where indicated. Experimental traces obtained from SPR sensograms for BG505 SOSIP binding to mAbs are also shown. SPR experiments are represented in black and best global fits are indicated in red. 1:1 Langmuir binding model and steady-state were used to calculate the association (ka) and dissociation (kd) rate constants.

**S5 Fig. Crystal structures of ElsE and Bess Fabs.**

The light and heavy chains are colored light and dark blue, respectively, with the ultralong CDR H3s highlighted in red. Two structures (ElsE8 and ElsE11) contain two Fabs in the crystallographic asymmetric unit, with both shown here. CDR H3 knob regions have very weak electron density in the Bess4 and ElsE8 mol1 structures that are not modeled. Extreme flexibility in the stalk regions is evident among these structures, for example when comparing the two Fab molecules in the asymmetric unit for ElsE11, where the knob domains are rotated approximately 114° from each other. **(A)** Fab Bess4, 2.8Å, **(B)** Fab Bess7, 2.1Å, **(C)** ElsE1, 1.81Å, **(D)** ElsE2,1.90Å, **(E)** ElsE5, 1.89Å, **(F)** ElsE6, 2.35Å, **(G)** ElsE7, 2.54Å, **(H)** ElsE8 mol1, 1.83Å, **(I)** ElsE8 mol2, 1.83Å, **(J)** ElsE9, 2.3Å, **(K)** ElsE11 mol1, 2.65Å, **(L)** ElsE11 mol2, 2.65Å. **(M)** Structural similarities of the Bess7 and PGT145 CDR H3 regions. The Loop region of the Bess7 knob folds into a 14 residues β-hairpin, with i-i+3 residues DEYA at the distal tip. The long CDR H3 of human anti-HIV Fab PGT145 has a similar type I β-turn at its tip, with i-i+3 residues NETysG. **(N)** Negative stain 3D reconstructions of Bess1, Bess2, Bess3 or Bess4 in complex with BG505 SOSIP and base-binding Fab RM20A3 (added for angular sampling).

**S6 Fig. Cryo-EM data processing and atomic modeling details of Bess4 in complex with BG505 SOSIP (PDB: 8TQ1). (A)** Representative 2D class averages. **(B)** Fourier Shell Correlation resolution estimation. **(C)** Angular distribution. **(D)** Local resolution estimation (Å) of the Bess4-BG505-RM20A3 cryo-EM dataset. **(E)** EM map density (contoured at 5σ) and modeled residues of CDRH3. **(F)** Overlay of CDRH3 from Bess4 and four human bnAbs, with BG505 SOSIP trimer shown as surface transparency. **(G)** Predicted hydrogen bonds and salt bridges between Bess4 and BG505 SOSIP. **(H)** Unsharpened and low contour (2σ) map of Bess4-BG505-RM20A3 with the Fv portion of the Bess4 crystal structure docked in.

**S1 Table. Neutralization ID_50_ titers are shown for IgG purified sera from Day 359 of all four cows. Geomean ID_50_ values and percent breadth are shown at the bottom of the graph. ID_50_ values are shown as 1/dilution.**

**S2 Table. Table of recovered heavy chains tested with native and universal light chains.**

**S3 Table. ElsE and Bess antibodies were tested for their ability to neutralize the 12-virus global panel. IC_50_ (µg/ml), IC_80_ (µg/ml), and MNP (%) are shown for all antibodies with a starting concentration of 50 µg/ml whose IC_50_ reach at least 50% neutralization. MNP= Maximum Neutralized Percentage.**

**S4 Table: X-ray data collection and refinement statistics for Bess and ElsE Fabs.**

**S5 Table. Cryo-EM data collection, refinement and validation statistics.**

**Movie S1: 3D variability analysis of cryo-EM data demonstrates movement of Bess4 mAb relative to BG505 SOSIP. Related to Figure 6**.

## References

1. Burton DR, Hangartner L. Broadly Neutralizing Antibodies to HIV and Their Role in Vaccine Design. Annu Rev Immunol. 2016;34:635–59. Epub 2016/05/12. doi: 10.1146/annurev-immunol-041015-055515. PubMed PMID: 27168247; PubMed Central PMCID: PMCPMC6034635.

2. Voss JE, Andrabi R, McCoy LE, de Val N, Fuller RP, Messmer T, et al. Elicitation of Neutralizing Antibodies Targeting the V2 Apex of the HIV Envelope Trimer in a Wild-Type Animal Model. Cell Rep. 2017;21(1):222–35. Epub 2017/10/06. doi: 10.1016/j.celrep.2017.09.024. PubMed PMID: 28978475; PubMed Central PMCID: PMCPMC5640805.

3. Pauthner M, Havenar-Daughton C, Sok D, Nkolola JP, Bastidas R, Boopathy AV, et al. Elicitation of Robust Tier 2 Neutralizing Antibody Responses in Nonhuman Primates by HIV Envelope Trimer Immunization Using Optimized Approaches. Immunity. 2017;46(6):1073–88 e6. Epub 2017/06/22. doi: 10.1016/j.immuni.2017.05.007. PubMed PMID: 28636956; PubMed Central PMCID: PMCPMC5483234.

4. Cottrell CA, van Schooten J, Bowman CA, Yuan M, Oyen D, Shin M, et al. Mapping the immunogenic landscape of near-native HIV-1 envelope trimers in non-human primates. PLoS Pathog. 2020;16(8):e1008753. Epub 2020/09/01. doi: 10.1371/journal.ppat.1008753. PubMed PMID: 32866207; PubMed Central PMCID: PMCPMC7485981.

5. Havenar-Daughton C, Lee JH, Crotty S. Tfh cells and HIV bnAbs, an immunodominance model of the HIV neutralizing antibody generation problem. Immunol Rev. 2017;275(1):49–61. Epub 2017/01/31. doi: 10.1111/imr.12512. PubMed PMID: 28133798.

6. Hu JK, Crampton JC, Cupo A, Ketas T, van Gils MJ, Sliepen K, et al. Murine Antibody Responses to Cleaved Soluble HIV-1 Envelope Trimers Are Highly Restricted in Specificity. J Virol. 2015;89(20):10383–98. Epub 2015/08/08. doi: 10.1128/JVI.01653-15. PubMed PMID: 26246566; PubMed Central PMCID: PMCPMC4580201.

7. Andrabi R, Voss JE, Liang CH, Briney B, McCoy LE, Wu CY, et al. Identification of Common Features in Prototype Broadly Neutralizing Antibodies to HIV Envelope V2 Apex to Facilitate Vaccine Design. Immunity. 2015;43(5):959–73. Epub 2015/11/21. doi: 10.1016/j.immuni.2015.10.014. PubMed PMID: 26588781; PubMed Central PMCID: PMCPMC4654981.

8. Gorman J, Soto C, Yang MM, Davenport TM, Guttman M, Bailer RT, et al. Structures of HIV-1 Env V1V2 with broadly neutralizing antibodies reveal commonalities that enable vaccine design. Nat Struct Mol Biol. 2016;23(1):81–90. Epub 2015/12/23. doi: 10.1038/nsmb.3144. PubMed PMID: 26689967; PubMed Central PMCID: PMCPMC4833398.

9. Doria-Rose NA, Bhiman JN, Roark RS, Schramm CA, Gorman J, Chuang GY, et al. New Member of the V1V2-Directed CAP256-VRC26 Lineage That Shows Increased Breadth and Exceptional Potency. J Virol. 2016;90(1):76–91. Epub 2015/10/16. doi: 10.1128/JVI.01791-15. PubMed PMID: 26468542; PubMed Central PMCID: PMCPMC4702551.

10. Doria-Rose NA, Schramm CA, Gorman J, Moore PL, Bhiman JN, DeKosky BJ, et al. Developmental pathway for potent V1V2-directed HIV-neutralizing antibodies. Nature. 2014;509(7498):55-62. Epub 2014/03/05. doi: 10.1038/nature13036. PubMed PMID: 24590074; PubMed Central PMCID: PMCPMC4395007.

11. Pancera M, McLellan JS, Wu X, Zhu J, Changela A, Schmidt SD, et al. Crystal structure of PG16 and chimeric dissection with somatically related PG9: structure-function analysis of two quaternary-specific antibodies that effectively neutralize HIV-1. J Virol. 2010;84(16):8098–110. Epub 2010/06/12. doi: 10.1128/JVI.00966-10. PubMed PMID: 20538861; PubMed Central PMCID: PMCPMC2916520.

12. Walker LM, Huber M, Doores KJ, Falkowska E, Pejchal R, Julien JP, et al. Broad neutralization coverage of HIV by multiple highly potent antibodies. Nature. 2011;477(7365):466-70. Epub 2011/08/19. doi: 10.1038/nature10373. PubMed PMID: 21849977; PubMed Central PMCID: PMCPMC3393110.

13. Briney BS, Willis JR, Crowe JE, Jr. Human peripheral blood antibodies with long HCDR3s are established primarily at original recombination using a limited subset of germline genes. PLoS One. 2012;7(5):e36750. Epub 2012/05/17. doi: 10.1371/journal.pone.0036750. PubMed PMID: 22590602; PubMed Central PMCID: PMCPMC3348910.

14. Shi B, Ma L, He X, Wang X, Wang P, Zhou L, et al. Comparative analysis of human and mouse immunoglobulin variable heavy regions from IMGT/LIGM-DB with IMGT/HighV-QUEST. Theor Biol Med Model. 2014;11:30. Epub 2014/07/06. doi: 10.1186/1742-4682-11-30. PubMed PMID: 24992938; PubMed Central PMCID: PMCPMC4085081.

15. Kodangattil S, Huard C, Ross C, Li J, Gao H, Mascioni A, et al. The functional repertoire of rabbit antibodies and antibody discovery via next-generation sequencing. MAbs. 2014;6(3):628–36. Epub 2014/02/01. doi: 10.4161/mabs.28059. PubMed PMID: 24481222; PubMed Central PMCID: PMCPMC4011907.

16. Lavinder JJ, Hoi KH, Reddy ST, Wine Y, Georgiou G. Systematic characterization and comparative analysis of the rabbit immunoglobulin repertoire. PLoS One. 2014;9(6):e101322. Epub 2014/07/01. doi: 10.1371/journal.pone.0101322. PubMed PMID: 24978027; PubMed Central PMCID: PMCPMC4076286.

17. Vigdorovich V, Oliver BG, Carbonetti S, Dambrauskas N, Lange MD, Yacoob C, et al. Repertoire comparison of the B-cell receptor-encoding loci in humans and rhesus macaques by next-generation sequencing. Clin Transl Immunology. 2016;5(7):e93. Epub 2016/08/16. doi: 10.1038/cti.2016.42. PubMed PMID: 27525066; PubMed Central PMCID: PMCPMC4973324.

18. Deiss TC, Vadnais M, Wang F, Chen PL, Torkamani A, Mwangi W, et al. Immunogenetic factors driving formation of ultralong VH CDR3 in Bos taurus antibodies. Cell Mol Immunol. 2019;16(1):53–64. Epub 2017/12/05. doi: 10.1038/cmi.2017.117. PubMed PMID: 29200193; PubMed Central PMCID: PMCPMC6318308.

19. Ma L, Qin T, Chu D, Cheng X, Wang J, Wang X, et al. Internal Duplications of DH, JH, and C Region Genes Create an Unusual IgH Gene Locus in Cattle. J Immunol. 2016;196(10):4358–66. Epub 2016/04/08. doi: 10.4049/jimmunol.1600158. PubMed PMID: 27053761.

20. Safonova Y, Shin SB, Kramer L, Reecy J, Watson CT, Smith TPL, et al. Variations in antibody repertoires correlate with vaccine responses. Genome Res. 2022;32(4):791–804. Epub 2022/04/02. doi: 10.1101/gr.276027.121. PubMed PMID: 35361626; PubMed Central PMCID: PMCPMC8997358.

21. Saini SS, Hein WR, Kaushik A. A single predominantly expressed polymorphic immunoglobulin VH gene family, related to mammalian group, I, clan, II, is identified in cattle. Mol Immunol. 1997;34(8-9):641–51. Epub 1997/06/01. doi: 10.1016/s0161-5890(97)00055-2. PubMed PMID: 9393967.

22. Wang F, Ekiert DC, Ahmad I, Yu W, Zhang Y, Bazirgan O, et al. Reshaping antibody diversity. Cell. 2013;153(6):1379-93. Epub 2013/06/12. doi: 10.1016/j.cell.2013.04.049. PubMed PMID: 23746848; PubMed Central PMCID: PMCPMC4007204.

23. Berens SJ, Wylie DE, Lopez OJ. Use of a single VH family and long CDR3s in the variable region of cattle Ig heavy chains. Int Immunol. 1997;9(1):189–99. Epub 1997/01/01. doi: 10.1093/intimm/9.1.189. PubMed PMID: 9043960.

24. Lopez O, Perez C, Wylie D. A single VH family and long CDR3s are the targets for hypermutation in bovine immunoglobulin heavy chains. Immunol Rev. 1998;162:55–66. Epub 1998/05/29. doi: 10.1111/j.1600-065x.1998.tb01429.x. PubMed PMID: 9602352.

25. Saini SS, Allore B, Jacobs RM, Kaushik A. Exceptionally long CDR3H region with multiple cysteine residues in functional bovine IgM antibodies. Eur J Immunol. 1999;29(8):2420–6. Epub 1999/08/24. doi: 10.1002/(SICI)1521-4141(199908)29:08<2420::AID-IMMU2420>3.0.CO;2-A. PubMed PMID: 10458755.

26. Saini SS, Kaushik A. Extensive CDR3H length heterogeneity exists in bovine foetal VDJ rearrangements. Scand J Immunol. 2002;55(2):140–8. Epub 2002/03/19. doi: 10.1046/j.1365-3083.2002.01028.x. PubMed PMID: 11896930.

27. Saini SS, Farrugia W, Ramsland PA, Kaushik AK. Bovine IgM antibodies with exceptionally long complementarity-determining region 3 of the heavy chain share unique structural properties conferring restricted VH + Vlambda pairings. Int Immunol. 2003;15(7):845–53. Epub 2003/06/17. doi: 10.1093/intimm/dxg083. PubMed PMID: 12807823.

28. de los Rios M, Criscitiello MF, Smider VV. Structural and genetic diversity in antibody repertoires from diverse species. Curr Opin Struct Biol. 2015;33:27–41. Epub 2015/07/21. doi: 10.1016/j.sbi.2015.06.002. PubMed PMID: 26188469; PubMed Central PMCID: PMCPMC7039331.

29. Zhao Y, Jackson SM, Aitken R. The bovine antibody repertoire. Dev Comp Immunol. 2006;30(1-2):175–86. Epub 2005/08/02. doi: 10.1016/j.dci.2005.06.012. PubMed PMID: 16054212.

30. Wu L, Oficjalska K, Lambert M, Fennell BJ, Darmanin-Sheehan A, Ni Shuilleabhain D, et al. Fundamental characteristics of the immunoglobulin VH repertoire of chickens in comparison with those of humans, mice, and camelids. J Immunol. 2012;188(1):322–33. Epub 2011/12/02. doi: 10.4049/jimmunol.1102466. PubMed PMID: 22131336.

31. Shojaei F, Saini SS, Kaushik AK. Unusually long germline DH genes contribute to large sized CDR3H in bovine antibodies. Mol Immunol. 2003;40(1):61–7. Epub 2003/08/12. doi: 10.1016/s0161-5890(03)00098-1. PubMed PMID: 12909131.

32. Smider BA, Smider VV. Formation of ultralong DH regions through genomic rearrangement. BMC Immunol. 2020;21(1):30. Epub 2020/06/04. doi: 10.1186/s12865-020-00359-8. PubMed PMID: 32487018; PubMed Central PMCID: PMCPMC7265228.

33. Jenkins GW, Safonova Y, Smider VV. Germline-Encoded Positional Cysteine Polymorphisms Enhance Diversity in Antibody Ultralong CDR H3 Regions. J Immunol. 2022;209(11):2141–8. Epub 2022/11/26. doi: 10.4049/jimmunol.2200455. PubMed PMID: 36426974; PubMed Central PMCID: PMCPMC9940733.

34. Huang R, Warner Jenkins G, Kim Y, Stanfield RL, Singh A, Martinez-Yamout M, et al. The smallest functional antibody fragment: Ultralong CDR H3 antibody knob regions potently neutralize SARS-CoV-2. Proc Natl Acad Sci U S A. 2023;120(39):e2303455120. Epub 2023/09/18. doi: 10.1073/pnas.2303455120. PubMed PMID: 37722054.

35. Stanfield RL, Berndsen ZT, Huang R, Sok D, Warner G, Torres JL, et al. Structural basis of broad HIV neutralization by a vaccine-induced cow antibody. Sci Adv. 2020;6(22):eaba0468. Epub 2020/06/11. doi: 10.1126/sciadv.aba0468. PubMed PMID: 32518821; PubMed Central PMCID: PMCPMC7253169.

36. Stanfield RL, Haakenson J, Deiss TC, Criscitiello MF, Wilson IA, Smider VV. The Unusual Genetics and Biochemistry of Bovine Immunoglobulins. Adv Immunol. 2018;137:135–64. Epub 2018/02/20. doi: 10.1016/bs.ai.2017.12.004. PubMed PMID: 29455846; PubMed Central PMCID: PMCPMC5935254.

37. Koti M, Kataeva G, Kaushik AK. Novel atypical nucleotide insertions specifically at VH-DH junction generate exceptionally long CDR3H in cattle antibodies. Mol Immunol. 2010;47(11-12):2119–28. Epub 2010/05/04. doi: 10.1016/j.molimm.2010.02.014. PubMed PMID: 20435350.

38. Liljavirta J, Niku M, Pessa-Morikawa T, Ekman A, Iivanainen A. Expansion of the preimmune antibody repertoire by junctional diversity in Bos taurus. PLoS One. 2014;9(6):e99808. Epub 2014/06/14. doi: 10.1371/journal.pone.0099808. PubMed PMID: 24926997; PubMed Central PMCID: PMCPMC4057420.

39. Liljavirta J, Ekman A, Knight JS, Pernthaner A, Iivanainen A, Niku M. Activation-induced cytidine deaminase (AID) is strongly expressed in the fetal bovine ileal Peyer’s patch and spleen and is associated with expansion of the primary antibody repertoire in the absence of exogenous antigens. Mucosal Immunol. 2013;6(5):942–9. Epub 2013/01/10. doi: 10.1038/mi.2012.132. PubMed PMID: 23299615.

40. Haakenson JK, Huang R, Smider VV. Diversity in the Cow Ultralong CDR H3 Antibody Repertoire. Front Immunol. 2018;9:1262. Epub 2018/06/20. doi: 10.3389/fimmu.2018.01262. PubMed PMID: 29915599; PubMed Central PMCID: PMCPMC5994613.

41. Stanfield RL, Wilson IA, Smider VV. Conservation and diversity in the ultralong third heavy-chain complementarity-determining region of bovine antibodies. Sci Immunol. 2016;1(1). Epub 2016/08/31. doi: 10.1126/sciimmunol.aaf7962. PubMed PMID: 27574710; PubMed Central PMCID: PMCPMC5000368.

42. Kramski M, Center RJ, Wheatley AK, Jacobson JC, Alexander MR, Rawlin G, et al. Hyperimmune bovine colostrum as a low-cost, large-scale source of antibodies with broad neutralizing activity for HIV-1 envelope with potential use in microbicides. Antimicrob Agents Chemother. 2012;56(8):4310–9. Epub 2012/06/06. doi: 10.1128/AAC.00453-12. PubMed PMID: 22664963; PubMed Central PMCID: PMCPMC3421555.

43. Heydarchi B, Center RJ, Gonelli C, Muller B, Mackenzie C, Khoury G, et al. Repeated Vaccination of Cows with HIV Env gp140 during Subsequent Pregnancies Elicits and Sustains an Enduring Strong Env-Binding and Neutralising Antibody Response. PLoS One. 2016;11(6):e0157353. Epub 2016/06/15. doi: 10.1371/journal.pone.0157353. PubMed PMID: 27300145; PubMed Central PMCID: PMCPMC4907510.

44. Heydarchi B, Center RJ, Bebbington J, Cuthbertson J, Gonelli C, Khoury G, et al. Trimeric gp120-specific bovine monoclonal antibodies require cysteine and aromatic residues in CDRH3 for high affinity binding to HIV Env. MAbs. 2017;9(3):550–66. Epub 2016/12/21. doi: 10.1080/19420862.2016.1270491. PubMed PMID: 27996375; PubMed Central PMCID: PMCPMC5384801.

45. Kramski M, Lichtfuss GF, Navis M, Isitman G, Wren L, Rawlin G, et al. Anti-HIV-1 antibody-dependent cellular cytotoxicity mediated by hyperimmune bovine colostrum IgG. Eur J Immunol. 2012;42(10):2771–81. Epub 2012/06/26. doi: 10.1002/eji.201242469. PubMed PMID: 22730083.

46. Sanders RW, Derking R, Cupo A, Julien JP, Yasmeen A, de Val N, et al. A next-generation cleaved, soluble HIV-1 Env trimer, BG505 SOSIP.664 gp140, expresses multiple epitopes for broadly neutralizing but not non-neutralizing antibodies. PLoS Pathog. 2013;9(9):e1003618. Epub 2013/09/27. doi: 10.1371/journal.ppat.1003618. PubMed PMID: 24068931; PubMed Central PMCID: PMCPMC3777863.

47. Sok D, Le KM, Vadnais M, Saye-Francisco KL, Jardine JG, Torres JL, et al. Rapid elicitation of broadly neutralizing antibodies to HIV by immunization in cows. Nature. 2017;548(7665):108-11. Epub 2017/07/21. doi: 10.1038/nature23301. PubMed PMID: 28726771; PubMed Central PMCID: PMCPMC5812458.

48. Heydarchi B, Fong DS, Gao H, Salazar-Quiroz NA, Edwards JM, Gonelli CA, et al. Broad and ultra-potent cross-clade neutralization of HIV-1 by a vaccine-induced CD4 binding site bovine antibody. Cell Rep Med. 2022;3(5):100635. Epub 2022/05/19. doi: 10.1016/j.xcrm.2022.100635. PubMed PMID: 35584627; PubMed Central PMCID: PMCPMC9133467.

49. Andrabi R, Pallesen J, Allen JD, Song G, Zhang J, de Val N, et al. The Chimpanzee SIV Envelope Trimer: Structure and Deployment as an HIV Vaccine Template. Cell Rep. 2019;27(8):2426–41 e6. Epub 2019/05/23. doi: 10.1016/j.celrep.2019.04.082. PubMed PMID: 31116986; PubMed Central PMCID: PMCPMC6533203.

50. Montefiori DC, Karnasuta C, Huang Y, Ahmed H, Gilbert P, de Souza MS, et al. Magnitude and breadth of the neutralizing antibody response in the RV144 and Vax003 HIV-1 vaccine efficacy trials. J Infect Dis. 2012;206(3):431–41. Epub 2012/05/29. doi: 10.1093/infdis/jis367. PubMed PMID: 22634875; PubMed Central PMCID: PMCPMC3392187.

51. deCamp A, Hraber P, Bailer RT, Seaman MS, Ochsenbauer C, Kappes J, et al. Global panel of HIV-1 Env reference strains for standardized assessments of vaccine-elicited neutralizing antibodies. J Virol. 2014;88(5):2489–507. Epub 2013/12/20. doi: 10.1128/JVI.02853-13. PubMed PMID: 24352443; PubMed Central PMCID: PMCPMC3958090.

52. Seaman MS, Janes H, Hawkins N, Grandpre LE, Devoy C, Giri A, et al. Tiered categorization of a diverse panel of HIV-1 Env pseudoviruses for assessment of neutralizing antibodies. J Virol. 2010;84(3):1439–52. Epub 2009/11/27. doi: 10.1128/JVI.02108-09. PubMed PMID: 19939925; PubMed Central PMCID: PMCPMC2812321.

53. Sok D, van Gils MJ, Pauthner M, Julien JP, Saye-Francisco KL, Hsueh J, et al. Recombinant HIV envelope trimer selects for quaternary-dependent antibodies targeting the trimer apex. Proc Natl Acad Sci U S A. 2014;111(49):17624–9. Epub 2014/11/26. doi: 10.1073/pnas.1415789111. PubMed PMID: 25422458; PubMed Central PMCID: PMCPMC4267403.

54. McCoy LE, Falkowska E, Doores KJ, Le K, Sok D, van Gils MJ, et al. Incomplete Neutralization and Deviation from Sigmoidal Neutralization Curves for HIV Broadly Neutralizing Monoclonal Antibodies. PLoS Pathog. 2015;11(8):e1005110. Epub 2015/08/13. doi: 10.1371/journal.ppat.1005110. PubMed PMID: 26267277; PubMed Central PMCID: PMCPMC4534392 following competing interests: EF, HS and TW are employed by the commercial companies, Abcam Burlingame, Crucell Holland B.V. and Monogram Biosciences Inc. respectively. This does not alter our adherence to all PLOS Pathogens policies on sharing data and materials.

55. Walker LM, Phogat SK, Chan-Hui PY, Wagner D, Phung P, Goss JL, et al. Broad and potent neutralizing antibodies from an African donor reveal a new HIV-1 vaccine target. Science. 2009;326(5950):285-9. Epub 2009/09/05. doi: 10.1126/science.1178746. PubMed PMID: 19729618; PubMed Central PMCID: PMCPMC3335270.

56. Honnen WJ, Krachmarov C, Kayman SC, Gorny MK, Zolla-Pazner S, Pinter A. Type-specific epitopes targeted by monoclonal antibodies with exceptionally potent neutralizing activities for selected strains of human immunodeficiency virus type 1 map to a common region of the V2 domain of gp120 and differ only at single positions from the clade B consensus sequence. J Virol. 2007;81(3):1424–32. Epub 2006/11/24. doi: 10.1128/JVI.02054-06. PubMed PMID: 17121806; PubMed Central PMCID: PMCPMC1797533.

57. Pinter A, Honnen WJ, D’Agostino P, Gorny MK, Zolla-Pazner S, Kayman SC. The C108g epitope in the V2 domain of gp120 functions as a potent neutralization target when introduced into envelope proteins derived from human immunodeficiency virus type 1 primary isolates. J Virol. 2005;79(11):6909–17. Epub 2005/05/14. doi: 10.1128/JVI.79.11.6909-6917.2005. PubMed PMID: 15890930; PubMed Central PMCID: PMCPMC1112130.

58. Doores KJ, Burton DR. Variable loop glycan dependency of the broad and potent HIV-1-neutralizing antibodies PG9 and PG16. J Virol. 2010;84(20):10510–21. Epub 2010/08/06. doi: 10.1128/JVI.00552-10. PubMed PMID: 20686044; PubMed Central PMCID: PMCPMC2950566.

59. Macpherson A, Laabei M, Ahdash Z, Graewert MA, Birtley JR, Schulze ME, et al. The allosteric modulation of complement C5 by knob domain peptides. Elife. 2021;10. Epub 2021/02/12. doi: 10.7554/eLife.63586. PubMed PMID: 33570492; PubMed Central PMCID: PMCPMC7972453.

60. Macpherson A, Scott-Tucker A, Spiliotopoulos A, Simpson C, Staniforth J, Hold A, et al. Isolation of antigen-specific, disulphide-rich knob domain peptides from bovine antibodies. PLoS Biol. 2020;18(9):e3000821. Epub 2020/09/05. doi: 10.1371/journal.pbio.3000821. PubMed PMID: 32886672; PubMed Central PMCID: PMCPMC7498065 following competing interests: All authors apart from J.v.d.E are current or previous employees of UCB-Celltech and may hold shares and/or stock options.

61. Adams R, Joyce C, Kuravskiy M, Harrison K, Ahdash Z, Balmforth M, et al. Serum albumin binding knob domains engineered within a V(H) framework III bispecific antibody format and as chimeric peptides. Front Immunol. 2023;14:1170357. Epub 2023/05/30. doi: 10.3389/fimmu.2023.1170357. PubMed PMID: 37251411; PubMed Central PMCID: PMCPMC10213618.

62. Hawkins A, Joyce C, Brady K, Hold A, Smith A, Knight M, et al. The proximity of the N- and C-termini of bovine knob domains enable engineering of target specificity into polypeptide chains. MAbs. 2022;14(1):2076295. Epub 2022/06/01. doi: 10.1080/19420862.2022.2076295. PubMed PMID: 35634719; PubMed Central PMCID: PMCPMC9154775.

63. Klewinghaus D, Pekar L, Arras P, Krah S, Valldorf B, Kolmar H, et al. Grabbing the Bull by Both Horns: Bovine Ultralong CDR-H3 Paratopes Enable Engineering of ‘Almost Natural’ Common Light Chain Bispecific Antibodies Suitable For Effector Cell Redirection. Front Immunol. 2021;12:801368. Epub 2022/01/29. doi: 10.3389/fimmu.2021.801368. PubMed PMID: 35087526; PubMed Central PMCID: PMCPMC8787767.

64. Pekar L, Klewinghaus D, Arras P, Carrara SC, Harwardt J, Krah S, et al. Milking the Cow: Cattle-Derived Chimeric Ultralong CDR-H3 Antibodies and Their Engineered CDR-H3-Only Knobbody Counterparts Targeting Epidermal Growth Factor Receptor Elicit Potent NK Cell-Mediated Cytotoxicity. Front Immunol. 2021;12:742418. Epub 2021/11/12. doi: 10.3389/fimmu.2021.742418. PubMed PMID: 34759924; PubMed Central PMCID: PMCPMC8573386.

65. Yanakieva D, Vollmer L, Evers A, Siegmund V, Arras P, Pekar L, et al. Cattle-derived knob paratopes grafted onto peripheral loops of the IgG1 Fc region enable the generation of a novel symmetric bispecific antibody format. Front Immunol. 2023;14:1238313. Epub 2023/11/09. doi: 10.3389/fimmu.2023.1238313. PubMed PMID: 37942319; PubMed Central PMCID: PMCPMC10628450.

66. Moyer TJ, Kato Y, Abraham W, Chang JYH, Kulp DW, Watson N, et al. Engineered immunogen binding to alum adjuvant enhances humoral immunity. Nat Med. 2020;26(3):430–40. Epub 2020/02/19. doi: 10.1038/s41591-020-0753-3. PubMed PMID: 32066977; PubMed Central PMCID: PMCPMC7069805.

67. Pugach P, Ozorowski G, Cupo A, Ringe R, Yasmeen A, de Val N, et al. A native-like SOSIP.664 trimer based on an HIV-1 subtype B env gene. J Virol. 2015;89(6):3380–95. Epub 2015/01/16. doi: 10.1128/JVI.03473-14. PubMed PMID: 25589637; PubMed Central PMCID: PMCPMC4337520.

68. Lefranc MP, Giudicelli V, Ginestoux C, Jabado-Michaloud J, Folch G, Bellahcene F, et al. IMGT, the international ImMunoGeneTics information system. Nucleic Acids Res. 2009;37(Database issue):D1006-12. Epub 2008/11/04. doi: 10.1093/nar/gkn838. PubMed PMID: 18978023; PubMed Central PMCID: PMCPMC2686541.

69. Otwinowski Z, Minor W. Processing of X-ray diffraction data collected in oscillation mode. Methods Enzymol. 1997;276:307–26. Epub 1997/01/01. doi: 10.1016/S0076-6879(97)76066-X. PubMed PMID: 27754618.

70. McCoy AJ, Grosse-Kunstleve RW, Adams PD, Winn MD, Storoni LC, Read RJ. Phaser crystallographic software. J Appl Crystallogr. 2007;40(Pt 4):658-74. Epub 2007/08/01. doi: 10.1107/S0021889807021206. PubMed PMID: 19461840; PubMed Central PMCID: PMCPMC2483472.

71. Emsley P, Lohkamp B, Scott WG, Cowtan K. Features and development of Coot. Acta Crystallogr D Biol Crystallogr. 2010;66(Pt 4):486–501. Epub 2010/04/13. doi: 10.1107/S0907444910007493. PubMed PMID: 20383002; PubMed Central PMCID: PMCPMC2852313.

72. Afonine PV, Grosse-Kunstleve RW, Echols N, Headd JJ, Moriarty NW, Mustyakimov M, et al. Towards automated crystallographic structure refinement with phenix.refine. Acta Crystallogr D Biol Crystallogr. 2012;68(Pt 4):352–67. Epub 2012/04/17. doi: 10.1107/S0907444912001308. PubMed PMID: 22505256; PubMed Central PMCID: PMCPMC3322595.

73. Berndsen ZT, Chakraborty S, Wang X, Cottrell CA, Torres JL, Diedrich JK, et al. Visualization of the HIV-1 Env glycan shield across scales. Proc Natl Acad Sci U S A. 2020;117(45):28014–25. Epub 2020/10/24. doi: 10.1073/pnas.2000260117. PubMed PMID: 33093196; PubMed Central PMCID: PMCPMC7668054.

74. Suloway C, Pulokas J, Fellmann D, Cheng A, Guerra F, Quispe J, et al. Automated molecular microscopy: the new Leginon system. J Struct Biol. 2005;151(1):41–60. Epub 2005/05/14. doi: 10.1016/j.jsb.2005.03.010. PubMed PMID: 15890530.

75. Zheng SQ, Palovcak E, Armache JP, Verba KA, Cheng Y, Agard DA. MotionCor2: anisotropic correction of beam-induced motion for improved cryo-electron microscopy. Nat Methods. 2017;14(4):331–2. Epub 2017/03/03. doi: 10.1038/nmeth.4193. PubMed PMID: 28250466; PubMed Central PMCID: PMCPMC5494038.

76. Punjani A, Rubinstein JL, Fleet DJ, Brubaker MA. cryoSPARC: algorithms for rapid unsupervised cryo-EM structure determination. Nat Methods. 2017;14(3):290–6. Epub 2017/02/07. doi: 10.1038/nmeth.4169. PubMed PMID: 28165473.

77. Pettersen EF, Goddard TD, Huang CC, Couch GS, Greenblatt DM, Meng EC, et al. UCSF Chimera--a visualization system for exploratory research and analysis. J Comput Chem. 2004;25(13):1605–12. Epub 2004/07/21. doi: 10.1002/jcc.20084. PubMed PMID: 15264254.

78. Casanal A, Lohkamp B, Emsley P. Current developments in Coot for macromolecular model building of Electron Cryo-microscopy and Crystallographic Data. Protein Sci. 2020;29(4):1069–78. Epub 2019/11/16. doi: 10.1002/pro.3791. PubMed PMID: 31730249; PubMed Central PMCID: PMCPMC7096722.

79. Conway P, Tyka MD, DiMaio F, Konerding DE, Baker D. Relaxation of backbone bond geometry improves protein energy landscape modeling. Protein Sci. 2014;23(1):47–55. Epub 2013/11/23. doi: 10.1002/pro.2389. PubMed PMID: 24265211; PubMed Central PMCID: PMCPMC3892298.

80. Afonine PV, Poon BK, Read RJ, Sobolev OV, Terwilliger TC, Urzhumtsev A, et al. Real-space refinement in PHENIX for cryo-EM and crystallography. Acta Crystallogr D Struct Biol. 2018;74(Pt 6):531–44. Epub 2018/06/07. doi: 10.1107/S2059798318006551. PubMed PMID: 29872004; PubMed Central PMCID: PMCPMC6096492.

